# FENNEC: Fine-Tuned Ensemble Neural Networks Accelerate Chemically Modified siRNA Screening and Design

**DOI:** 10.64898/2026.06.13.732049

**Authors:** Alexander Larsen, Johannes Braun, Rachapun Rotrattanadumrong, Philipp Berninger, Dimitar Yonchev, Julien Gagneur, Daniel Butnaru, Annalisa Marsico

**Affiliations:** Computational Health Center (CHC), Helmholtz Zentrum Muenchen, Ingolstaedter Landstrasse 1, Oberschleissheim, 80939, Germany; Pharma Research and Early Development, Roche Innovation Center Basel, Grenzacherstrasse 124, Basel, 4070, Switzerland; TUM School of Computation, Information and Technology (CIT), Technical University of Munich, Boltzmannstrasse 3, Garching bei Muenchen, 85748, Germany; Cluster for Nucleic Acid Sciences and Technologies – NUCLEATE, Feodor-Lynen-Str. 25, D-81377 München

**Keywords:** siRNA, Neural Networks, Convolution, LLMs, RNAi, RNA interference, RNA therapeutics, Chemically modified siRNA

## Abstract

Small interfering RNAs (siRNAs) are a clinically validated therapeutic modality, but designing potent chemically modified siRNAs remains costly and iterative due to limited public data. Computational prediction of siRNA efficacy can accelerate rational design and preclinical development. However, despite the importance of chemical modifications in therapeutic performance, current machine-learning methods either do not account for this or generalize poorly beyond their training data. Here, we present FENNEC (Fine-Tuned Ensemble of Neural Networks for siRNA Efficiency Characterization), a machine-learning framework for predicting chemically modified siRNA activity. We curate the largest publicly available patent-derived dataset of modified siRNAs from 42 patents using OCR-based table extraction and stringent quality filtering. FENNEC combines temporal convolutional networks with thermodynamic features, experimental covariates, and RNA foundation-model embeddings to capture sequence- and transcript-level determinants of efficacy. We show that foundation-model embeddings encode transferable biological information, particularly benefiting prediction in data-scarce settings. FENNEC achieved robust gene- and siRNA-level performance, including independent validation on a novel AHSA1-targeting dataset, and consistently outperformed existing approaches. Model interpretation recovers modification-specific effects, while in silico perturbation analyses demonstrate its utility for both prediction and design optimization of chemically modified siRNAs. Together, these results establish FENNEC as a chemically aware framework for data-driven siRNA design. We have FENNEC which will be publicly available for non-commercial use at https://github.com/marsico-lab/fennec.

## INTRODUCTION

Small interfering RNAs (siRNAs) have emerged as a transformative therapeutic modality, capable of targeting, in principle, any disease-associated genes with high specificity [1, 2]. These short, double-stranded RNA duplexes (typically 19–23 bp) harness the endogenous RNA-induced silencing complex (RISC) to mediate sequence-specific mRNA degradation [3]. However, clinical efficacy relies, among other things, on extensive chemical modifications to enhance stability, reduce immunogenicity, and promote uptake. While modifications like 2’-O-methyl, 2’-fluoro, and phosphorothioate backbones are essential for drug properties such as stability, enhanced target affinity, reduced off-targeting and nuclease degradation resistance [4, 5], they introduce complex, non-linear effects on silencing potency that simple sequence-based rules cannot capture. Davis et al., recently established a design framework showing that the efficacy of fully chemically modified siRNAs depends on the precise positioning of modifications and the accessibility of the target mRNA structure[6]. However, designing highly potent, chemically modified siRNAs remains a costly, iterative process driven largely by trial-and-error [7].

Computational prediction offers a path to accelerate this cycle, but current methods present several limitations. Classical machine learning models such as random forests, linear regression, and support vector machines (SVM), with engineered features such as GC content, minimum free energy, and positional information trained on unmodified sequence efficacy data [8, 9, 10, 11, 12, 13] fail to generalize to the chemically modified regime. Recent deep learning approaches employing convolutional neural networks, Multi-layer perceptrons, and transformers to predict siRNA efficacy [14, 15, 16, 17], while powerful, lack access to large-scale, diverse datasets of modified siRNAs [18]. Furthermore, public datasets often lack key experimental covariates, such as transfection method, cell line, and siRNA concentration, which are significant confounders in knockdown prediction. This data scarcity and heterogeneity have prevented the development of robust, chemistry-aware predictors.

Recent RNA foundation models, including RNA-FM, RiNALMo, and Orthrus, have demonstrated strong predictive performance across diverse RNA tasks such as structure inference, RNA type/family classification, RNA–protein interaction prediction, functional property prediction, and predicting the effects of steric-blocking oligonucleotides (SBOs) on gene expression and splicing—highlighting their potential for broad application in RNA therapeutic design [19, 20, 21, 22, 23]. OligoFormer employs the RNA-FM language model to generate context-aware embeddings of siRNAs and target sequences, capturing features beyond simple sequence composition. However, it is trained on unmodified datasets, is limited to a guide length of 19 bp, and does not support chemically modified nucleotides. Other models which are chemically aware, such as cm-siRNA-pred and Martinelli’s models both from 2024 [14, 15], are likely limited by the dataset they were trained on, siRNAMod, which despite representing a substantial compilation effort, contains predominantly small, heterogeneous experimental batches with limited coverage of clinically relevant modification patterns. We hypothesize that integrating these pre-trained embeddings can capture complex biological priors—specifically local mRNA accessibility profiles and RNA-binding protein (RBP) occupancy probabilities—that are difficult to encode manually using standard thermodynamic features (e.g. MFE), thereby enhancing efficacy prediction in the modified regime.

To bridge this gap, we present FENNEC (Fine-Tuned Ensemble of Neural Networks for siRNA Efficiency Characterization), a deep learning model designed to predict siRNA knockdown (KD) efficacy that learns joint representations of chemically modified siRNAs, their target mRNAs, and relevant experimental metadata, trained on a large-scale curated dataset of patent-extracted measurements for modified siRNA sequences. FENNEC treats siRNA design as a multi-modal learning problem. The model architecture (Fig.1, described in Methods) combines three information streams: (1) a one-hot encoding of the guide and passenger strands, capturing local sequence-chemistry patterns, i.e. the exact placement of *x* possible chemical modifications; (2) thermodynamic and reaction-condition metadata, allowing the model to adjust predictions based on assay context; (3) and latent representations from pre-trained RNA language models that contribute target information. The inputs are processed through an ensemble from 10 seeds of temporal convolutional networks (TCNs) and multi-headed attention layers before converging into a final Multi-layer Perceptron (MLP) to output the predicted mRNA remaining percentage upon knockdown — a widely used empirical proxy for siRNA silencing activity that reflects the net outcome of RISC loading, target cleavage, and assay conditions, without explicitly modelling any individual mechanistic step. This hybrid approach allows FENNEC to learn both the “grammar” of chemical modifications and the underlying biological constraints of the target mRNA.

**Figure 1.**
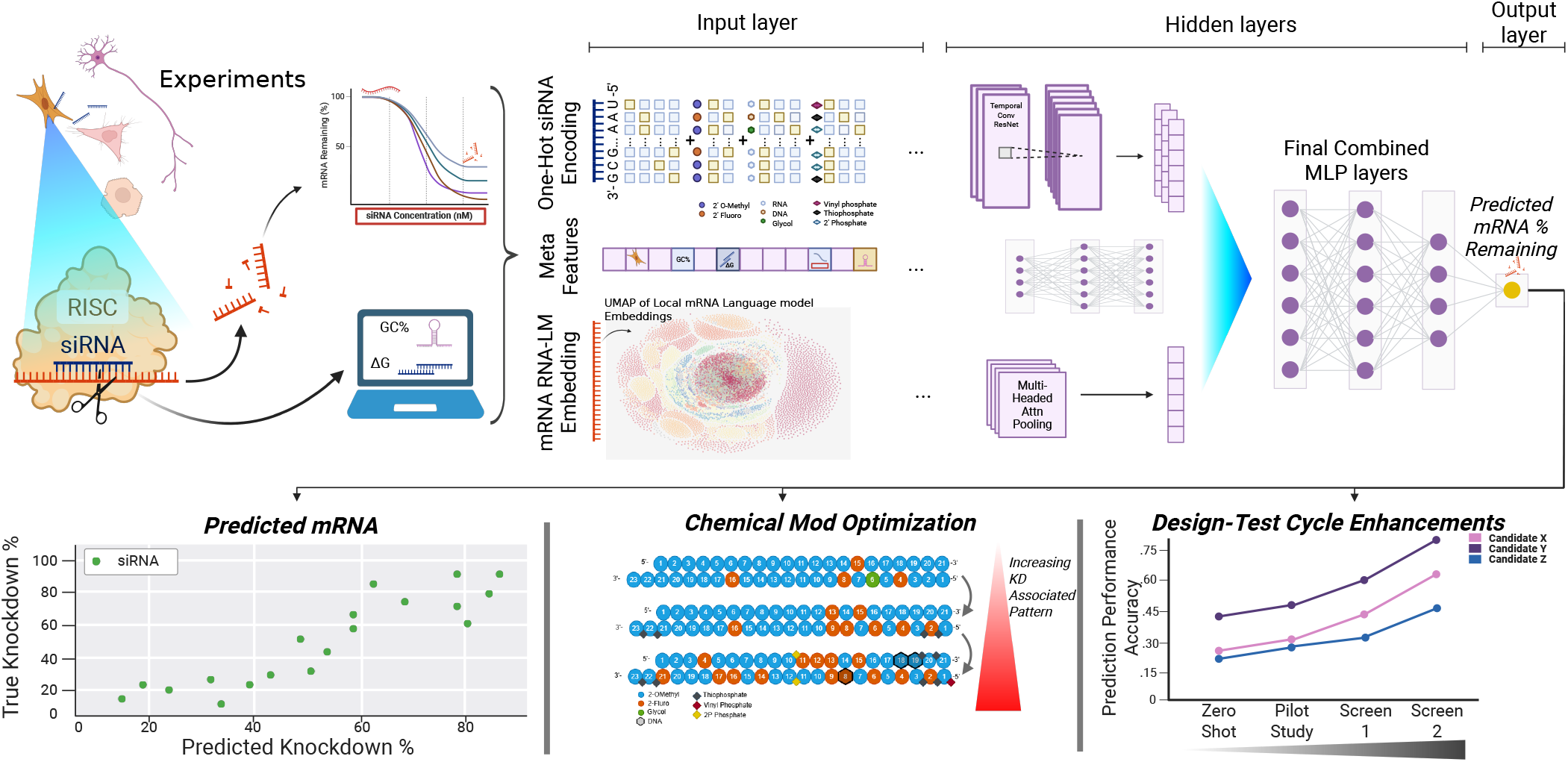
Graphical Overview of the FENNEC Model and Its Applications. Using Experimental data, calculated features, and Foundational Model Embeddings on 100bp of the 5’UTR start, 100bp flanking window of the siRNA target, and 100bp of the 3’UTR start separately, we employ the FENNEC architecture to train a model capable of predicting mRNA Knock Down (KD), which can be used as an oracle for siRNA chemical optimization, hotspot prediction, and we show it can be easily tailored to new low N datasets via our data efficiency studies.

The main contributions of this work are:

- We curated a large machine-readable dataset of chemically modified siRNAs from 42 patents spanning 35 gene targets and integrated these with existing unmodified data, linking sequence and chemical modification information to assay metadata across 76 gene targets, 16 cell types, 11 concentrations, approximately 13,500 unique siRNA guide–passenger sequences, and 12 clinically-relevant distinct chemical modification types (excluding unmodified RNA). To our knowledge, this represents the broadest combination of target, cell type and concentration diversity and chemical modification coverage reported to date.
- We demonstrated that FENNEC accurately predicts siRNA knockdown and consistently outperforms existing approaches across multiple benchmark model categories.
- In a Design-test Cycle experiment we show that FENNEC can be rapidly applied to new, smaller datasets/targets, significantly improving accuracy from “zero-shot” through subsequent screening phases. In this setting, we highlight the added value of RNA language-model embeddings in data-scarce scenarios, which reflect the limited sample sizes typical of pilot experiments in pharmaceutical research
- We validate FENNEC robustness by applying it to novel previously unseen target sequences. Experimental results, testing 94 siRNAs targeting the AHSA1 transcript, confirm that FENNEC maintains high predictive accuracy in these new biological contexts.
- Finally, we demonstrate that FENNEC can be applied as predictive “oracle” for optimal chemical modification patterns (shown by the blue and orange patterns) to maximize silencing.

To our knowledge, we established FENNEC as a chemistry-aware and interpretable framework capable of predicting siRNA efficacy across diverse experimental conditions.

## MATERIALS AND METHODS

### Data Preprocessing

We searched Google Patents for siRNA-related patents filed by Alnylam Pharmaceuticals (WO patent office, January 2013 to September 2025) using the query ‘RNAi methods and compositions’, identifying 153 candidate patents. Publicly available unmodified siRNA datasets from Huesken et al., Katoh et al., and Ichihara et al.[8, 9, 10] were included to supplement coverage of unmodified sequence space.

Patent tables were extracted from PDF images using PaddleOCR[24] with the PP-StructureV3 pipeline, employing RT-DETR-L wired for table-cell detection, SLANeXt wired for table structure recognition, and en PP-OCRv4 mobile rec for text recognition (minimum detector side length 3000 px; det thresh = 0.15, box thresh = 0.4, unclip ratio 1.5–2.0). Following extraction, sequence tables were merged with their corresponding knockdown readout tables, linking each siRNA to its measured mRNA remaining percentage per cell line and concentration. For patents where OCR recovery was incomplete, an internal cross-referencing tool (PatentMiner) was used as a supplement or replacement — most notably for the chemically diverse APP dataset. Target mRNA sequences were retrieved from NCBI using the Entrez command-line tool: `efetch -db nucleotide -id` *<*`accession`*>* `-format fasta`[25].

A general overview of the data curation pipeline, from patent scraping to quality control, is shown in Fig. 2A. A complete list of all patents used for training and testing, with corresponding gene IDs, table titles, and datapoint counts, is provided in Table S3. The listed counts reflect only successfully extracted and quality-filtered datapoints; many patents contained additional measurements that failed extraction or were removed during downstream filtering.

**Figure 2.**
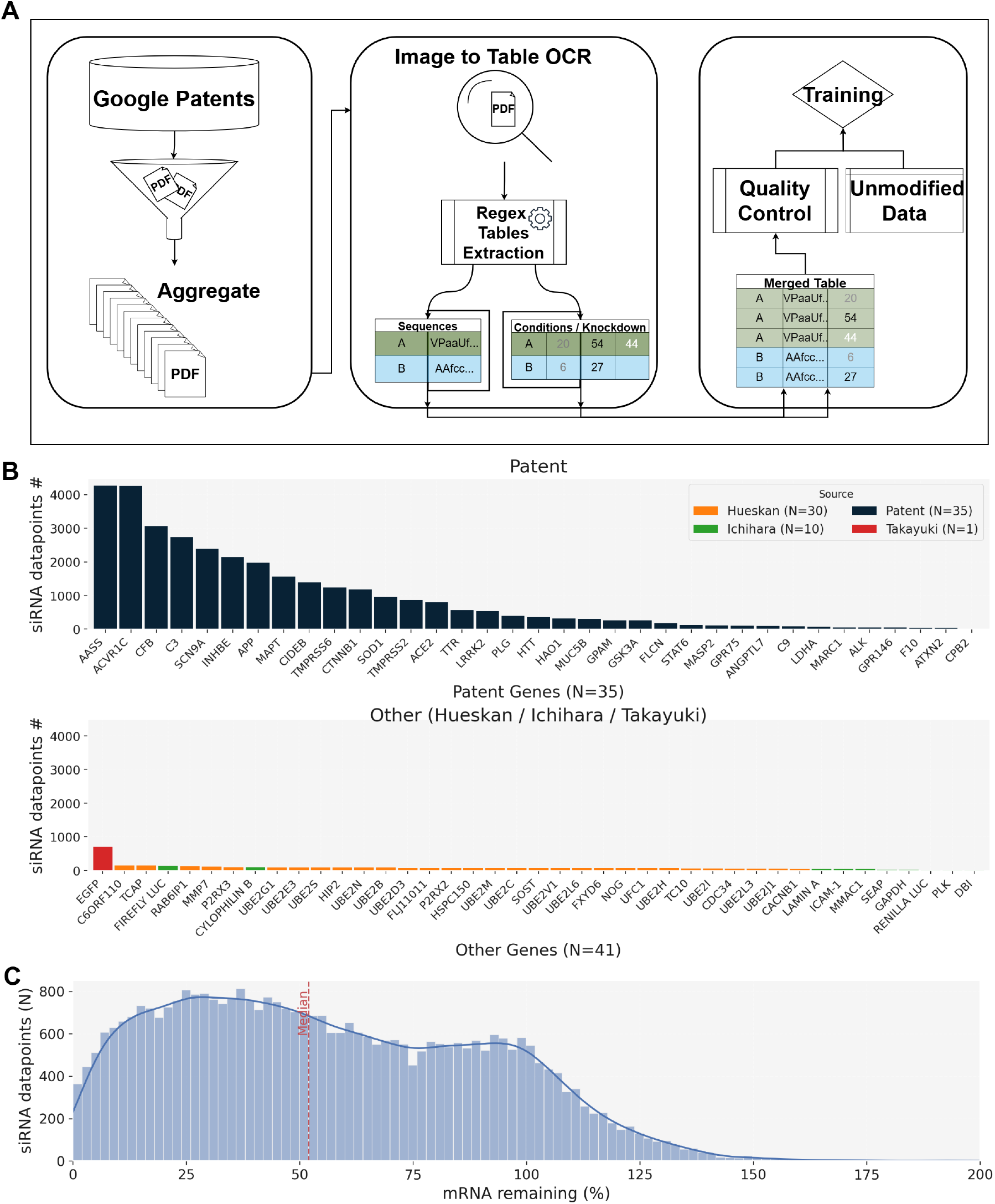
Dataset overview. (A) A summarization of the scraping procedure. (B) Distribution of curated chemically modified siRNA datapoints per target gene for the patents and the older sets (some are removed based on alignment to target). (C) distribution of efficiency mRNA remaining % used as *Y*_true_.

### Quality Control

The initial dataset was filtered using standard experimental design and data quality principles. Patents were retained only if they demonstrated high-quality text extraction, defined as the ability to accurately merge table cell content and maintain 95% column datatype purity. We required retrieved siRNAs to have a minimum length of 16 nucleotides for both forward and reverse strands and to explicitly contain modified nucleotides exceeding 20% of the sequence.

Data points with unclear or unlisted target gene assignments or lacking an associated transcript were excluded. To ensure correct guide-passenger pairing, we applied homology filters by transforming chemically modified siRNAs into simple ATGC format. Using the Python package `edlib`, we calculated the edit distance between the guide and the reverse complement of the passenger strand; pairs with an edit distance greater than six, including a common 2nt overhang of the guide, were excluded to filter out potential OCR row-shift errors and the length of the guide-passenger pair was required to be 2bp or 0bp - the most common pattern observed in patents. The reverse complement of each guide was similarly required to align to the extracted mRNA strand with an edit distance of at most 6. Finally, compounds with unclear concentration usage, unknown chemical modifications, or fewer than 10 associated chemical modifications were removed.

After applying all filters, the final dataset comprised 36,394 datapoints from 42 patents covering 35 patent-derived gene targets. Combined with the unmodified sequence datasets, the corpus spans 76 unique target genes, 13,647 unique guide-passenger sequence combinations, and 1,134 distinct chemical modification patterns. The chemical vocabulary includes RNA, DNA, and glycol nucleic acid (GNA) backbone types; 2’-O-methyl, 2’-fluoro, and 2’-hexadecyl sugar modifications; and backbone linkers and conjugates including phosphorothioate, vinylphosphonate, 2’-phosphate, and GalNAc (L96). Measurements span 16 cell types, 11 concentration levels, and multiple transfection methods and durations, preserving the experimental heterogeneity of the original patent assays.

### Model Architecture

*Primary Model Design* FENNEC’s architecture comprises three complementary modules. The first encodes the siRNA sequence and chemical modification pattern (Fig. S5) using a Residual Temporal Convolutional Network (TCN). Specifically, the encoder consists of a ResNet with asymmetric left-padding to enforce causality, ensuring that each sequence position attends only to itself and preceding positions. Three parallel 1D convolutional kernels with widths of 3, 5, and 7, and corresponding dilation rates of 1, 2, and 4, are applied within independent TCN blocks with residual skip connections. The resulting feature maps are concatenated along the sequence dimension and projected through a 128-dimensional multilayer perceptron (MLP). The second module processes thermodynamic meta-features (described below) using a separate three-layer MLP with 32-dimensional hidden layers. The third module encodes the RNA target using RNA foundation model embeddings, which are aggregated through a multi-head attention pooling layer [26, 27] with 4 heads, followed by a 2-layer 128-dimensional MLP. The final FENNEC prediction is the average output of an ensemble of ten independently seeded model instances, forming a regression over mRNA knockdown percentage clipped to the range (0–200).

#### *Features and Encodings* The model accepts three primary input streams

1. Guide siRNA/Chemistry/Passenger information encoded in a combination of one-hot encodings. For all positions, the following were encoded: nucleotide identity (A, T, U, G, C, I [inosine]), nucleic acid type (DNA, RNA, Glycol nucleic acid [GNA]), 2’-sugar modifications (e.g., 2’-F, 2’-OMe, 2’-hexadecyl), and phosphate linker types (e.g., vinylphosphonate, 2’-phosphate, standard phosphate, phosphorothioate). These encodings were concatenated along the sequence length and padded to a length of 25. The same was done for the passenger strand, and the two 25×*D*_*one*−*hotfeatures*_ sequences were concatenated.
2. Thermodynamic features partially retrieved from ViennaRNA, biopython, Primer3, including GC content, a custom Δ*G* distribution bias score, and predicted mRNA minimum free energy (MFE)[28, 29, 30]. Meta-features such as cell line donor, log-transformed siRNA concentration, transfection duration, and transfection method were also included[31].
3. LLM embeddings from RNA-FM, RiNALMo, and Orthrus, each computed on 100 bp windows of the target mRNA. We evaluated the influence of different transcript regions on model predictions by comparing RNA foundation model embeddings derived from distinct sequence contexts. Specifically, embeddings were generated from (i) the target region centered on the siRNA binding site (padded with *N* s where necessary), (ii) the first 100 bp of the 5’ UTR, (iii) the first 100 bp of the 3’ UTR immediately downstream of the stop codon, or (iv) the concatenation of all three regions into a single embedding. UTR boundaries were extracted from GenBank records retrieved via NCBI. Representations were taken from the 12th layer for RNA-FM and the standard pooled representation for RiNALMo and Orthrus [20, 21, 22].

### Model training and evaluation

#### Training Setup

For all experiments, we conducted 10-fold cross-validation across 10 random initialization seeds, employing a gene-based splitting strategy to ensure no overlap of target genes or base siRNAs between training and test sets. This means that for CV1, no siRNA sequence from any training genes would occur in the test set. By chance, some cell lines and concentrations may have been in only one split as a consequence. We also evaluated FENNEC using a “base siRNA” split, a data partitioning strategy commonly adopted in previous studies trained on unmodified siRNA datasets. Under this scheme, siRNAs targeting the same gene are likely to appear in both the training and test sets; however, individual guide–passenger sequence pairs are exclusive to a single split, ensuring that no exact siRNA is observed during training. This setting therefore evaluates generalization to unseen siRNAs within previously encountered gene targets, rather than to entirely novel targets. To prevent overfitting and optimize training duration, an inner 80/20 split was utilized to determine the ideal number of epochs for early stopping; this fixed epoch count was then used to train on the full fold before evaluating on the held-out outer loop test set for each of the 10 CV holdouts. While classical models were tuned via Optuna, the final configuration of FENNEC was optimized through small incremental improvements and a grid search to explore the number of TCN blocks, the convolution filter size used, the heads for the multi-headed attention pooling, and depth of Linear layers for the meta branch[32]. For training, we utilized the AdamW optimizer with a cosine annealing schedule, featuring a learning rate of 1 × 10^−3^, a batch size of 128, and a weight decay of 0.005. All models were trained using MSE loss; Huber loss was evaluated during hyperparameter tuning but abandoned due to equivalent performance.

### Data Efficiency and Transfer Learning Simulations

To evaluate FENNEC’s capacity to generalize and adapt to novel biological targets and unseen chemical spaces under data-scarce scenarios, we conducted simulated data efficiency (low-*N*) design cycles using two independent target genes held out from the primary training phase: *ACVR1C* (from patent WO-2025064660-A2) and *JAK1* (from patent WO-2024256707A1).

The simulations were structured as progressive data-incorporation experiments to mimic sequential stages of a pharmaceutical screening campaign:

1. **Target-Context Low-***N* **Adaptation (*ACVR1C*):** The *ACVR1C* dataset (*N* = 1,435) was utilized to evaluate target generalization on a transcript featuring an unusually long 3’ UTR. We simulated a baseline “zero-shot” deployment by excluding all *ACVR1C* data from the initial model weights. Subsequently, we progressively introduced increasing fractions of target-specific data (ranging from 0% to 80% in increments) into the training partition, evaluating model performance on the remaining held-out validation fraction. To quantify the specific contribution of structural language priors in low-sample regimes, these data-scaling trajectories were executed in parallel using FENNEC variants trained both with and without foundational RNA language model embeddings (Orthrus and RNA-FM). These models were evaluated across three independent concentration tiers (0.1 nM, 1 nM, and 10 nM).
2. **Out-of-Distribution Chemistry Adaptation (*JAK1*):** The *JAK1* dataset (*N* = 191) was selected to assess chemical pattern transfer learning, characterized by an out-of-distribution alternating 2’-F/2’-O-Me “checkerboard” modification architecture that was completely absent from the primary training corpus. Mirroring the *ACVR1C* workflow, the model was subjected to incremental fine-tuning steps, scaling up to an 80% training allocation (*N* = 153).

For all efficiency coordinates, the predictive accuracy was rigorously monitored using continuous rank-ordering tracking (Spearman rank correlation coefficient) alongside top-quartile binary classification performance (Area Under the Receiver Operating Characteristic, AUROC) to evaluate the model’s calibration stability as a function of sample size expansion.

### Zero-shot Benchmark Models

To demonstrate FENNEC’s added value, we benchmarked FENNEC against multiple external models. The following methods were selected for benchmarking. RNAxs [33] is a rule-based tool that scores target site accessibility via RNA secondary structure prediction and three established design criteria (strand asymmetry, self-folding energy, free-end unpaired nucleotides); it was selected for its historical significance and its shared use of the ViennaRNA package, though it was not trained on chemically modified siRNAs. OligoFormer [19] is a transformer-based model combining a CNN-based siRNA encoder, an RNA-FM language model for target context, and a thermodynamic module; its multi-branch architecture closely mirrors FENNEC’s design philosophy, making it a natural reference point for zero-shot performance on chemically modified sequences, despite being trained only on unmodified data.

Cm-siRPred [14] and the Monopoli-RF web server [6, 34] were selected as, to our knowledge, the only published methods that natively handle chemically modified siRNAs. Cm-siRPred employs multi-view cross-attention and a 3D-CNN operating on one-hot sequences, molecular fingerprints, and physicochemical properties. Monopoli is a regressor trained on chemically modified siRNA sequences using sequence and structural features and is accessible via a public web server. Both represent the most directly comparable baselines to FENNEC, albeit trained on considerably smaller datasets.

FENNEC was trained with early stopping using standarding training methods excluding JAK1 and APP. Due to the specific input constraints of each tool, the following preprocessing steps were applied to enable valid comparisons:

#### RNAXS

The standard RNAXS [33] implementation limits inputs to 19-mers. We modified the source Perl script to accept variable guide lengths, enabling processing of our full dataset.

#### Monopoli

Predictions were obtained via the public web server on December 20th, 2025 from http://sirna-frontend.s3-website.us-east-2.amazonaws.com/search. A subset of sequences were not processed by the server and were consequently excluded from the comparison.

#### OligoFormer

We evaluated two model variants [19]. For *OligoFormer (0-19bp)*, which is intolerant to mismatches, we truncated guide sequences to 19bp and accounted for the dataset’s normalization of 5’ nucleotides to U. The *OligoFormer (1-20bp)* variant was one-nt window over but not to the end of the 23mer.

#### cm-siRNA-pred

Using the cm-siRNA-pred [14] servers API on December 23rd, 2025, we mapped our chemical vocabulary to the model’s supported list. Two approximations were required: (1) linker modifications were excluded as they were not supported, and (2) GNA modifications were replaced with Serinol, the closest structural analog available in their library.

These adaptations were necessary to enable a zero-shot comparison.

### Classical Benchmark Models

To complement these external benchmarks and account for the inherent difficulties in cross-platform comparisons, we also evaluated classical machine learning models trained directly on our curated data. These included simple regression via ElasticNet, Ridge, Lasso, SVR, Random Forest, and XGBoost with hyperparameters tuned via Optuna [32] were evaluated via 10-fold cross validation. For these tests, overlapping target sequences were removed. Features for these models included a flattened positional one hot encoded information of the modified siRNA as well as the same engineered “thermo” features used in the top performing version of FENNEC shown in TableS1.

### AHSA1 Experimental Validation

#### siRNA Selection for Experimental Validation

We experimentally validated FENNEC’s siRNA potency predictions using AHSA1, an oncogenic regulator of HSP90 and a potential cancer biomarker and therapeutic target that is commonly used in siRNA knockdown studies. Candidate siRNAs targeting AHSA1 were selected using an internal prioritization algorithm applying human-specificity and seed-toxicity filters, yielding 94 sequences stratified across the predicted activity range to ensure broad coverage of model-predicted efficacy.

#### Cell Culture and siRNA Transfection

U251 cells (CVCL 0021) were maintained in RPMI 1640 media (Gibco, 61870) supplemented with 10% Fetal Bovine Serum (FBS) (Corning, 35-010-CF) and 1% Penicillin-Streptomycin (Pen-Strep) (Gibco, 15140). Cells were cultured at 37^°^C in a humidified atmosphere with 5% CO_2_. For siRNA transfection, U251 cells were reverse transfected using Lipofectamine RNAiMAX (ThermoFisher Scientific 13778150). A half-log dose response curve was generated with final siRNA concentrations ranging from 20 nM to 0.00064 nM and *IC*_50_ values, corresponding to the concentration of the siRNA that produces 50% of the maximum inhibition of the target, were computed by fitting those curves. siRNAs were acoustically dispensed into 384-well plates (Corning, 3764) using an Echo 550 (Beckman Coulter) acoustic liquid handler. A complex of 10 µl of Opti-MEM (Gibco, 51985-026) and 0.3 µl of RNAiMAX per well was added to the dispensed siRNAs. U251 cells were detached using 0.25% Trypsin-EDTA (Gibco, 25200-056) and 6000 cells in 40 µL of complete cell culture medium were added per well to the lipoplexes. After 24 hours of incubation, total RNA was isolated using the Echolution cell culture RNA kit (Bioecho Life Sciences, 011-314-008) following the manufacturer’s protocol. AHSA1 and HPRT1 mRNA quantification was performed using probe-based qPCR assays (IDT, AHSA1: Hs.PT.58.4598253, HPRT1: Hs.PT.58v.45621572) and *TaqPath*^*TM*^ 1-Step Multiplex Master Mix (ThermoFisher Scientific, A28527) on a QuantStudio 7 Pro (ThermoFisher Scientific) real-time qPCR thermal cycler.

Percentage of AHSA1 mRNA remaining was calculated for each siRNA and concentration by normalizing AHSA1 *Ct* values to HPRT1, a non-targeting housekeeping control gene, using the ΔΔ*Ct* method, and expressing the result relative to a non-targeting control transfection. This yielded a residual mRNA measurement directly comparable to FENNEC’s predicted output (mRNA remaining %). FENNEC predictions were compared against these normalized experimental measurements independently at each of the 10 tested concentrations (0.0006–20 nM). Concordance was quantified using Spearman rank correlation between predicted and measured residual mRNA, assessing the model’s ability to correctly rank siRNA potency. Linear regression of measured residual mRNA on predicted values was used to compute *R*^2^ as a measure of quantitative agreement, with regression statistics reported per concentration (Table S2). To assess whether FENNEC captures intrinsic siRNA potency independent of concentration, predicted residual mRNA was additionally correlated with absolute *IC*_50_ values derived from four-parameter logistic fits to the full dose–response curve for each siRNA. To benchmark FENNEC against competing methods (OligoFormer, Cm-siRPred) on this experimentally generated dataset, siRNAs were binarized into “highly active” (top quartile of lowest residual mRNA) versus the remainder at the concentration showing the strongest predictive relationship (6.33 nM), and performance was evaluated using precision–recall area under the curve (AUPRC) and top-k hit-rate analysis (Fig. 7D).

### Feature Attribution via Integrated Gradients

To interpret model predictions and quantify the contribution of individual sequence motifs and chemical modifications, we employed Integrated Gradients (IG) [35], a path-integrated gradient attribution method. IG satisfies the axiom of completeness, ensuring that the sum of feature attributions equals the difference between the model’s output for the input and a baseline reference.

Attributions were computed for the ensemble of models composing FENNEC using the Captum v0.7.0[36]. For each input sample *x* in the hold-out test set, we calculated attributions separately for two input modalities: the one-hot encoded siRNA sequence (*X*_seq_) and chemical modification modality using the python package Captum v0.7.0[36]. To isolate the effect of a specific modality, we integrated gradients along a linear path from a zero-baseline (*x*^*′*^ = 0) to the input *x* in 128 steps, while holding the other two input modalities constant at their true values. This formulation ensures that the relevance score for the *i*-th feature, *IG*_*i*_(*x*), reflects its contribution within the specific biochemical context of the molecule:

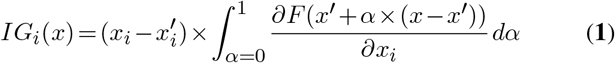

where *F* represents the regression model output (e.g., mRNA remaining). The integral was approximated using a Riemann sum. To compare feature sensitivity across metadata inputs with different scales (e.g., binary indicators vs. concentration), we normalized the mean integrated gradient scores by the global mean absolute value of each feature plus a small epsilon (*ϵ* = 1e-5), yielding a scale-invariant measure of feature potency. Final feature importance for a given position or channel was derived from the magnitude of these calculated attributions.

### Analysis of target transcript properties in embedding space

To determine what biophysical information the RNA language-model embeddings encode, we correlated their principal components with computed target-transcript features, such as gene length, RiboNN-predicted translation efficiency [37], exon-junction distance, and flanking GC content, among others. To derive target information, patent-derived siRNA records were loaded from the entire dataset and de-duplicated at the guide-passenger sequence level (retaining one entry per unique sequence pair, independent of experimental conditions, since transcript embeddings depend only on the mRNA sequence). Gene symbols were harmonized where necessary, and reported target mRNA sequences were curated against a manually aggregated reference FASTA. For target sequences shorter than 120 nucleotides, the reported sequence was converted to DNA alphabet and matched against reference transcripts after replacement of ambiguous R bases with N; when a match was found, the full reference transcript was substituted, otherwise the original reported target sequence was retained. Transcript-derived features were then computed from the corrected mRNA sequences, including melting temperature, GC fraction, polarity, GC skew, linguistic complexity (k = 4), predicted unpaired-sequence percentage, and Vienna minimum free energy. To annotate target-site context, each siRNA was aligned to gene-specific GenBank records using edlib in semi-global mode, and the best alignment was used to estimate the minimum distance to the nearest exon boundary and to classify the target site as 5’ untranslated region, coding sequence, or 3’ untranslated region. These annotations were used exclusively for embedding-space interpretation and were not included as model input features. Gene-level translational-efficiency annotations were calculated from RiboNN predictions and merged with the transcript metadata[37, 38].

Embedding analyses were performed using the FENNEC preprocessing framework. Transcript embedding tensors were generated with the Orthrus four-track triple representation using a slice length of 100 nucleotides for each of the 5’ UTR start, siRNA centered, and 3’UTR start; additional exploratory one-hot embeddings were also generated in parallel analyses. Principal component analysis (PCA) was applied to the embedding matrix, retaining 30 components for downstream analysis. Associations between PCA coordinates and transcript features were quantified using Spearman rank correlation and non-parametric mutual information. Features were partitioned into gene-level static and slice-level dynamic categories by testing within-gene variability; features with at most one unique value in at least 95% of genes were treated as static, whereas all remaining numeric features were treated as dynamic. Static-feature associations were summarized after collapsing observations to the per-gene median, whereas dynamic-feature associations were computed at the slice level. Correlation and mutual-information matrices were hierarchically ordered using average-linkage clustering with Euclidean distance. For visualization of local structure, UMAP was applied to the minimum number of leading principal components required to explain ≈85% of the total variance, using cosine distance and a fixed random seed.

### Chemical Architecture Optimization via Greedy Beam Search

Next, we tested whether FENNEC could serve as a predictive oracle for siRNA chemical optimization under controlled experimental conditions, where siRNA concentration, cell line donor, siRNA sequence (AUGC composition), and transfection method are kept constant, allowing us to isolate and quantify the contribution of chemical modifications alone.

The space of possible chemical modification patterns for a single siRNA duplex is combinatorially vast: with up to 12 modification types available at each of about 20 positions across guide and passenger strands, exhaustive evaluation of all possible chemistry for a single sequence is computationally intractable. To address this, we implemented an iterative greedy beam search algorithm operating on the FENNEC fitness landscape. A greedy beam search addresses this by treating chemical modification design as a guided local search problem: rather than evaluating every possible pattern, the algorithm iteratively proposes small, local perturbations to a starting design (single-position modification swaps), evaluates each proposal using a fitness oracle, and retains only the most promising candidates to seed the next round of exploration. This mirrors an experimentalist’s iterative optimization process, but executes it *in silico* across many candidate branches using quantitative guidance. The beam (a fixed-width set of retained top candidates at each round) allows exploration of multiple promising paths in parallel rather than committing to a single trajectory.

Our implementation explores local mutational neighborhoods to identify high-potency design trajectories, and adapts the beam search framework benchmarked in NucleoBench by Shor et al. [39].

#### Algorithm and Search Parameters

The optimization process is initialized using a parent siRNA sequence, denoted with Hamming distance *H* = 0 relative to itself (Figure S9), exhibiting suboptimal knockdown activity. The algorithm executes a directed “modification walk” through *I* discrete iterations (rounds; in our experiments *I* = 5). In each round *i*, a mutational neighborhood is generated by systematically substituting every possible chemical modification at every position of the double-stranded siRNA duplex relative to each candidate retained from the previous round, producing a set of single-edit candidate sequences at Hamming distance *i* from the original parent. Each candidate is scored using the FENNEC model (the oracle), and only the top *k* unique candidates (the beam size) are propagated to round *i*+1 (Figure S9).

To balance search depth with computational efficiency, we employed a variable beam width (*k*). We used a beam width of *k* = 50 was utilized sequences with expansive searchable areas, that is those with 1000+ possible combinations based on the existing base siRNA’s positional chemical substitution space. For restricted search spaces, with a smaller positional chemical substitution space i.e. of 6 unique modification patterns with base siRNA *S*, we only saw modifications in positions 2, 6, 8, and 14, which were one of three mods: 2^*′*^OMe, 2^*′*^F, or DNA for a total of 3^4^; the width was adjusted (e.g., *k* = 3) to ensure the algorithm traversed no more than 25% of the total possible modification combinations. At the conclusion of each iteration, only the top *k* unique candidates were propagated to the subsequent round. Adaptive *k* aligns with NucleoBench’s finding that fixed hyperparameters do not generalize across tasks, and should instead be calibrated to problem scale.

Figure S9 illustrates this process schematically for *k* = 2 (the real design used larger beam widths, as described above): starting from a parent siRNA at *H* = 0, each iteration expands every current beam member into its full single-position mutational neighborhood (candidates shown in blue/discarded candidates marked with a red ×), the FENNEC oracle scores all candidates, and only the top *k* performers are retained to seed the subsequent round. After *I* rounds, this process yields optimized candidate sequences at Hamming distance *H* = *I* from the original parent, representing the endpoint of the guided modification walk (green, Figure S9).

#### Calculation of the Enrichment Factor

To quantify the specific design utility of the FENNEC oracle, we calculated a **Recovery Enrichment Factor (EF)**, defined as the ratio of valid industrial matches recovered by the informed search versus a compute-based random baseline:

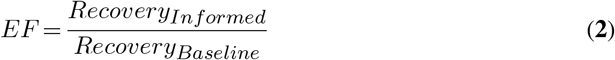

The baseline expected recovery was established through a stochastic search mirroring the architecture of the informed greedy beam. In this null model, the algorithm executes the same number of iterations and maintains an identical beam width (*k*); however, rather than prioritizing candidates via the FENNEC ensemble score, *k* variants are selected stochastically at each iteration for propagation to the subsequent round. This approach ensures that the calculated enrichment factor accounts for the local connectivity of the search space and reflects the true marginal benefit of model-guided prioritization over random neighborhood exploration.

#### Design Space Constraints: Closed vs

*Open World* Beyond simply finding a high-scoring pattern, we wanted to know how much of FENNEC’s design guidance reflects previously validated industrial chemistry versus genuinely novel combinations. To distinguish these, we ran the greedy beam search under two different constraints on which modifications are allowed at each position:

- **Closed-World (Local Cohort):** At each position, the search may only choose among modifications that were experimentally observed at that same position, for that same base sequence, within its own patent-derived cohort. This restricts exploration to a “known” design space and tests whether FENNEC can correctly re-prioritize modification patterns that have real industrial precedent
- **Open-World (Cohort-Pool):** At each position, the search may draw from the full pool of observed modifcations across all the study sequences (six in this case, totaling *N* = 43 unique position-chemistry combinations), regardless of whether that specific modification was ever tested at that specific position or in that specific sequence context. This mode allows for the exploration of novel, out-of-distribution patterns to characterize potential “design headroom.”

#### Heuristic Ranking and Uncertainty Quantitation

To ensure the selection of robust candidates and mitigate the impact of model outliers, we introduced another modification to the greedy beam search. Variants explored by the search were ranked using a lower-bound efficacy metric derived from the model ensemble, Rather than the FENNEC score itself. For each candidate sequence, the predicted mRNA remaining percentage was calculated as:

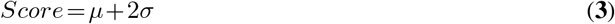

where *µ* represents the ensembled mean of predicted mRNA knockdown across the 10 random initialization seeds, and *σ* represents the standard deviation. This heuristic prioritization allows the algorithm to favor candidates that exhibit both high predicted potency and high ensembled consensus, effectively penalizing regions of high epistemic uncertainty in the fitness landscape.

## RESULTS

### Dataset Overview

The curation and filtering pipeline described in Methods yielded a dataset of 36,394 datapoints spanning 76 unique gene targets, 13,647 guide-passenger sequence combinations, and 1,134 distinct chemical modification patterns (Fig.2A–B). The composition of experimental covariates — transfection method, siRNA concentration, cell line, and evaluation time — is summarised in Fig.2C, and the distribution of mRNA remaining percentages and modification frequencies is shown in Fig.2D–E.

### FENNEC achieves state-of-the-art performance in Benchmark Studies

We evaluated FENNEC’s performance in the prediction of siRNA efficacy via mRNA remaining percent as a regression task of continuous range computing the true vs predicted spearman correlation, as well as in a binary classification task 50-50 split to evaluate the ROC-AUC. Model performance was evaluated using ten-fold cross-validation (see Methods) under two data-splitting strategies: by base siRNA (defined as sequences with identical nucleotide composition but distinct chemical modification patterns) and by target gene, the latter to assess potential target-specific biases. We further conducted an ablation study to quantify the contribution of each input modality and architectural component to the model performance, including a systematic comparison of different LLM-derived embeddings (RNA-FM, RiNALMo and Orthrus) of the target RNA. Finally, we benchmarked FENNEC against classical machine learning baselines trained on handcrafted features (see Methods).

The best model configuration (FENNEC + Orthrus Embeddings) achieves a Spearman-R of 0.52 and an AUC of 0.71 when split by gene, improving to a Spearman-R of 0.76 and an AUC of 0.80 when split by “base siRNA” (Fig.3A). The gap between these metrics separates the model’s ability to optimize chemistry on a known lead (0.76) from its ability to identify novel therapeutic candidates on unseen genes (0.52). All ablation studies were conducted on the gene split to strictly evaluate performance on novel gene targets. These comparisons (Fig. 3B) highlight the contribution of each architectural and engineered feature. Interestingly, ablating chemical modification features results in a 28% drop in performance (Spearman-R), compared to 20% and 18% decreases when removing metadata features and LLM-derived representations, respectively, highlighting the critical importance of chemical modification information for model predictions. The optimal model utilizes Orthrus embeddings (sequence length 100) with multi-headed attention pooling, the thermodynamic features listed in Supplementary TableS1, and a Residual TCN architecture. This result also suggests that Orthrus may capture a more informative and functionally relevant representation of the RNA target than the alternative language model, potentially due to its biologically informed contrastive loss. Using an ensemble approach—averaging predictions across 10 FENNEC runs with different random seeds—further improves performance, increasing both Spearman correlation and AUROC by up to 5% (Fig.3C-D). To validate the superiority of deep learning models, we benchmarked the ensembled version of FENNEC against five classical machine learning models Fig.3C-D using the same dataset and engineered features (see Methods). The best-performing classical model, XGBoostRegressor, achieved a Spearman-R of 0.42—inferior to standard versions of FENNEC, including many ablated variants with less performance driving features.

**Figure 3.**
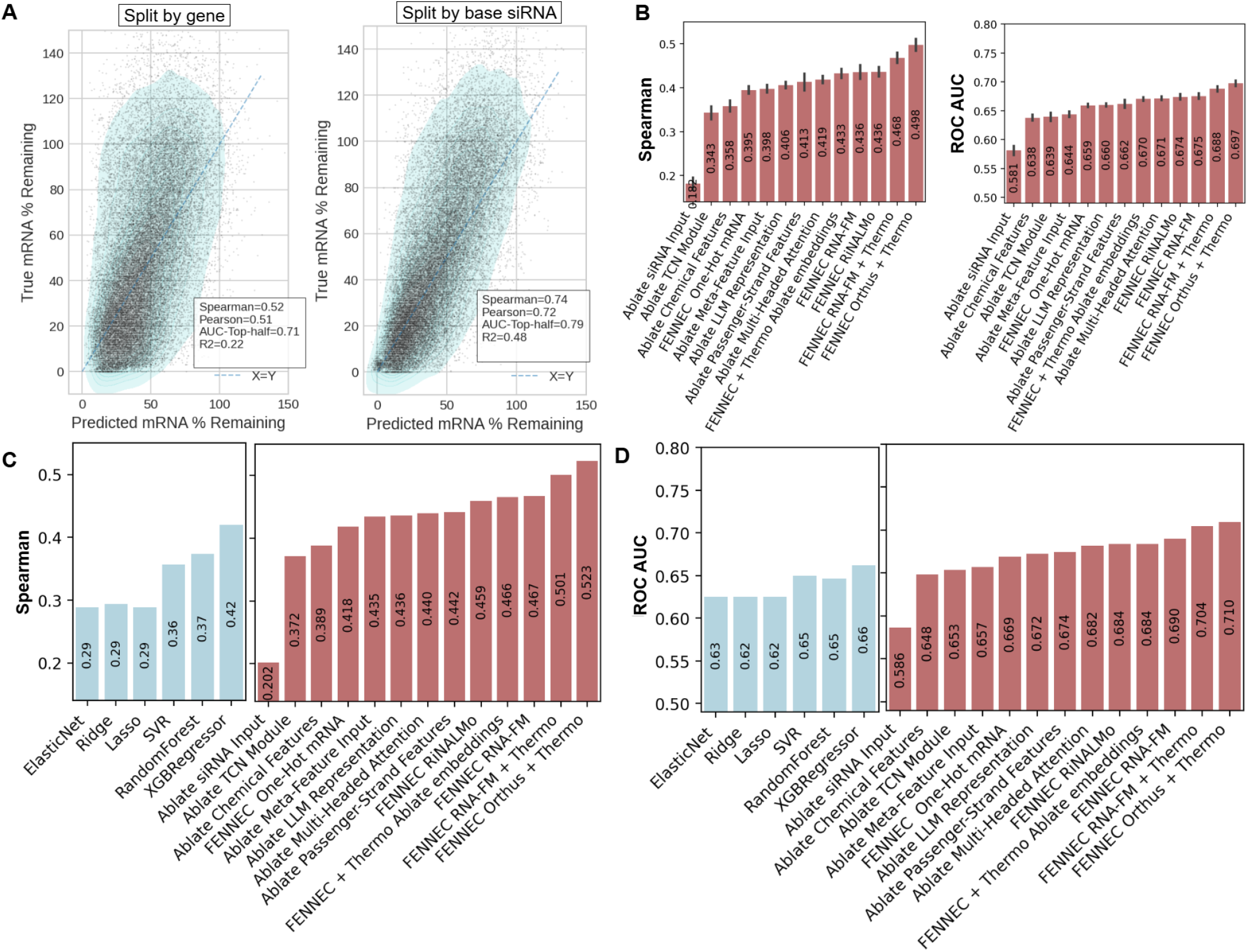
Model Performance: (A) Scatterplot by splitting strategy and the statistics associated with each. (B) 10 seeds per ablation model + alternate FM embeddings to test benefit of each architectural / data-stream. Error bars in Fig. represent 95% confidence intervals (CI) based on seed replicates.(C) Spearman correlation comparing the standard model baselines - left compared to the ensembled prediction of the 10 seeds - right. (D) AUC of the same models as a binary classification task separating the top 50% most effective compared to the bottom 50%.

### Data Efficiency: Boosting Performance on Novel Targets

We investigated whether FENNEC can predict and improve siRNA efficacy on new data, including novel gene targets or chemical modification patterns not seen during initial training, particularly in low-data settings. This setup mimics a typical experimental design cycle in pharmaceutical research, where models must generalize from a limited initial pool of siRNA sequences available for the novel target, with performance evolving as more data becomes available. We therefore assessed how predictive accuracy scales with increasing training set size, and whether specific components—such as RNA language model–derived representations—provide a distinct advantage in low-N regimes.

#### Atypical Target

*ACVR1C* We first conducted an efficiency study on a novel target absent from FENNEC’s training data—the ACVR1C gene from a recently published patent WO-2025064660-A2, where the pretrained model initially failed to predict siRNA efficacy (Spearman ≈0.05, AUROC ≈0.5, Fig.4A-B). ACVR1C presents an atypical prediction challenge: it harbours an exceptionally long 3’ UTR dense with regulatory elements reflecting its broad expression and pleiotropic signalling roles, making target-site accessibility highly context-dependent and poorly approximated by standard thermodynamic features alone. To address this, we progressively incorporated increasing fractions of target-specific data into the training set and evaluated performance at each stage. This allowed us to (1) determine the minimal data required to achieve satisfactory predictive performance, and (2) assess the impact of RNA language model (LM)–derived representations, which we hypothesised would be particularly informative given the gene’s complex 3’ UTR architecture.

FENNEC showed strong gains in low-data regimes, reaching ≈80% of its full Spearman performance and ≈90% of its AUROC with only 20% of the data ((Fig.4A-B), highlighting its utility in data-scarce settings such as early-stage experimental campaigns. Notably, in these low-N scenarios, performance was substantially improved by incorporating RNA target representations from the Orthrus LM, an effect not observed with RNA-FM and that diminished as training data increased. This underscores the value of learning functional RNA representations, particularly when limited data is available, in improving siRNA knockdown prediction.

Fig. 4A–B shows results at the 10 nM concentration; all three tested concentrations (0.1 nM, 1 nM, and 10 nM) show a similar pattern (Fig. S10).

**Figure 4.**
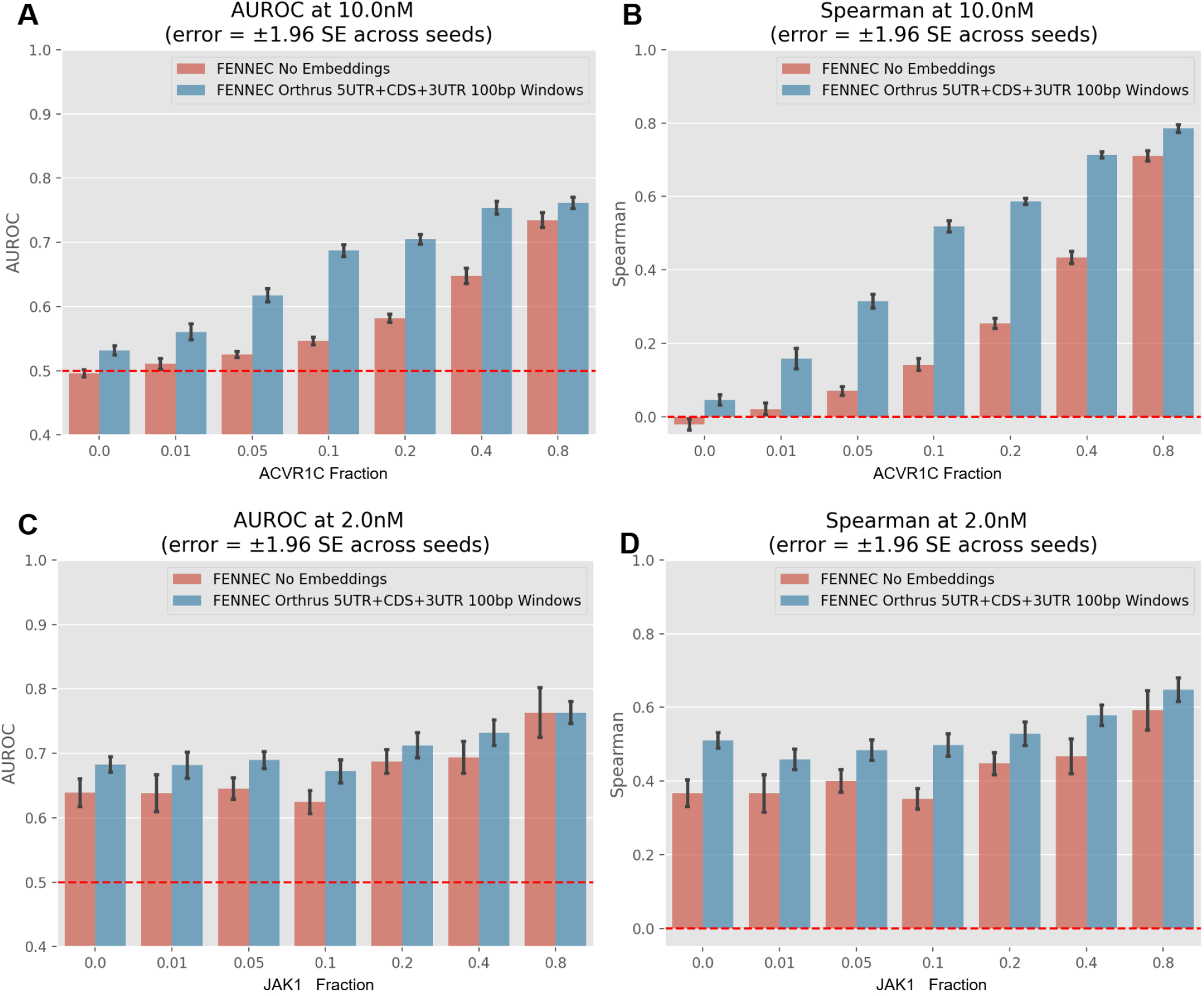
Data efficiency effects. (A-B) Barplots of FENNEC with and without foundation model embeddings (observed vs predicted knockdown) across increasing ACVR1C (N=1435) gene fraction mixed into training (0–80%), illustrating calibration effects on AUROC and Spearman. (C-D) Top-half AUROC and Spearman difference for JAK1 (N=191) - Alternative chemical pattern set with increasing JAK1 N in training.

#### Assessing Data Efficiency on Alternative Chemistry

*Data - the JAK1 case study* Second, we performed a similar data efficiency study to evaluate FENNEC’s ability to generalize to a new target, JAK1, absent from the original training set, featuring a set of N=191 sequences exhibiting an out-of-distribution chemical modification pattern, namely the checkerboard 2’-F/2’-O-Me design (alternating 2’-O-Me and 2’-F modifications). As shown in Fig. 4C– D, FENNEC achieved reasonable zero-shot performance (Spearman *r* = 0.506, AUROC = 0.68) and improved to a mean Spearman *r* = 0.67 with 80% of the data (*N* = 153) incorporated into training. Notably, incorporating RNA language model embeddings provided no meaningful advantage for JAK1, in contrast to the ACVR1C case. This contrast is informative: when the primary bottleneck is recognising an unfamiliar chemical modification pattern, the explicit chemistry encoding is sufficient and FM-derived target context adds little — whereas FM embeddings become valuable when the bottleneck is target-sequence generalisation to a biologically unusual transcript. FENNEC’s performance improving steadily as increasing fractions of new data are incorporated further demonstrates its ability to generalise to unseen chemical designs from other research groups.

### Zero-Shot Independent Test Set *In Silico* Benchmarking

Two zero-shot benchmarks were conducted to evaluate FENNEC against current state-of-the-art methods, including deep learning approaches (e.g., Cm-siRPred and OligoFormer) as well as more traditional models, such as RNAxs and the Monopoli-RF web server. These methods differ in their support for chemically modified siRNAs: RNAxs and OligoFormer were developed and trained only on datasets containing unmodified siRNAs, whereas Cm-siRPred and the Monopoli-RF web server were trained on datasets that include chemically modified siRNAs reported in the literature (see the Materials and Methods section).

The first benchmark used a curated chemically diverse subset of the Amyloid Precursor protein (APP) dataset from patent WO2020132227A2 consisting of 343 chemically modified siRNA sequences targeting various areas with a diverse set of chemical patterns not present in our training data with APP ablated. Unique in primarily its chemical diversity, it allows to evaluate how models with and without chemical modifications predict on such datasets. As shown in Table 1, on the APP dataset, FENNEC largely outperformed all benchmarked methods, achieving a Spearman correlation of 0.427, with RNAxs emerging as the closest competitor. OligoFormer and the remaining methods showed more limited generalization to the chemical modification pattern present in this dataset. The second benchmark focused on a JAK1 gene subset from patent ID WO2024256707A1 with a unique chemical patterns from a separate company, where FENNEC again demonstrated strong performance with a Spearman correlation of 0.506, outperforming all competitors, as detailed in Table 2.

**Table 1.** Zero-shot benchmark APP Chemically Diverse Subset.

| Model | Evaluated $N$ | Spearman | Pearson | AUC 25% | PR 25% |
| --- | --- | --- | --- | --- | --- |
| **FENNEC** | **343/343** | **0.427** | **0.402** | **0.674** | **0.384** |
| RNAxs Vienna | 343/343 | 0.297 | 0.271 | 0.620 | 0.328 |
| cm-siRNA-pred | 343/343 | 0.188 | 0.188 | 0.581 | 0.296 |
| Oligoformer_1_20<sup>†</sup> | 339/343 | 0.092 | 0.111 | 0.509 | 0.254 |
| Oligoformer_0_19<sup>†</sup> | 340/343 | 0.078 | 0.082 | 0.537 | 0.543 |

<sup>†</sup> Oligoformer primarily accepts 19bp inputs; indices 0\_19 and 1\_20 represent the two window positions tested.

**Table 2.**
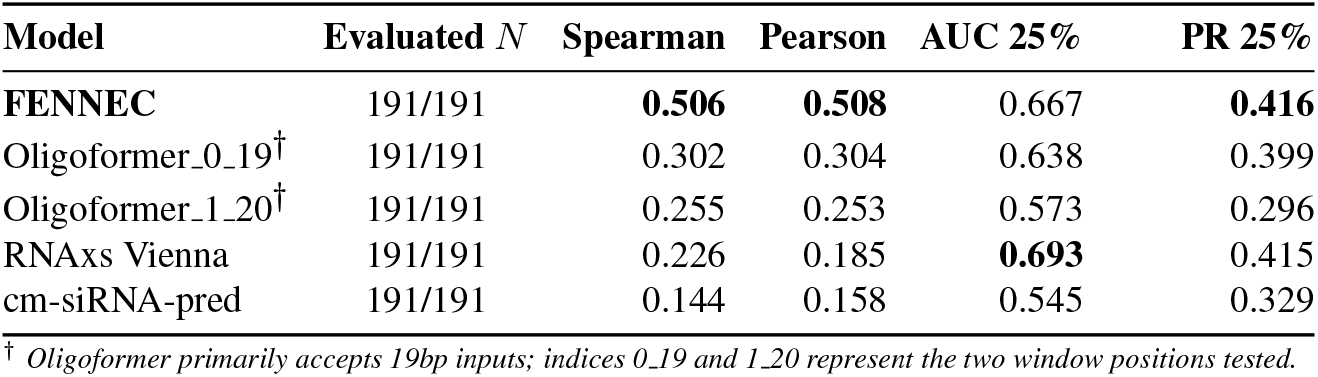
Zero-shot benchmark JAK1 unique chemical pattern subset.

An important consideration when interpreting these results is that, although we made the appropriate modifications to use all methods fairly for this inference task (see Methods), not all methods were compatible with the full dataset: the column labeled *Evaluated N* reports the percentage of the dataset successfully processed by each method, reflecting either input format and length incompatibilities or failures of the online prediction portal. Additionally, the predictions attributed to Monopoli should be interpreted with caution: no standalone implementation is publicly available, results were obtained through the web server under its published terms of use, and we cannot fully verify that the server was used under conditions optimal for our dataset. All in all, these results highlight FENNEC’s ability to generalize effectively to unseen chemical modifications, underscoring its potential utility in guiding siRNA design across diverse therapeutic targets.

### Model Interpretation

*Sequence Attribution* Drivers of model performance from the integrated gradients [35] calculated using python package Captum v0.7.0[36] are summarized in Fig.5. Sequence composition, shown in three panels Fig.5A-C shows AU preference at position 1, reduced cytosine content across positions 2–7 (seed), and higher GC tolerance in the 3’ end of the guide. Cytosine enrichment in the seed region appears to hurt knockdown slightly more than guanine and position 14 seems to additionally be hurt by higher G/A/C content compared to surrounding areas.

#### Chemical Attribution

Panels A-B of Fig.5 (guide and passenger) show strong associations at positions 1-7, 14, and 16, frequently involving GNA, DNA swap, hexadecyl, 2’-O-methyl (2’-OMe), and 2’-fluoro (2’-F) modifications, with individual positions showing a preference against G/C identity associated with reduced knockdown. Additional associations include a slightly positive associated vinylphosphonate at the guide 5’ end and a mild association for phosphorothioates near terminal positions of both strands with a more harmful effect of the phosphorothioates at the 5’ of the guide.

#### Reaction/Meta Attribution

Meta-features, comprising both reaction context variables and engineered thermodynamic descriptors, provide a measurable improvement in predictive performance. Compared with the meta-feature-ablated variant, the full FENNEC model achieves an increase of approximately 0.1 in Spearman correlation and 0.05 in accuracy (Fig. 5B). Feature importance analysis indicates that reaction context variables contribute most strongly to these gains, with transfection method and cell line emerging as the most influential predictors. In particular, free uptake is generally associated with reduced siRNA performance, whereas experiments performed in Neuro2a cells tend to show higher performance. By contrast, engineered thermodynamic descriptors, although occasionally informative, contribute less consistently than the reaction context variables.

**Figure 5.**
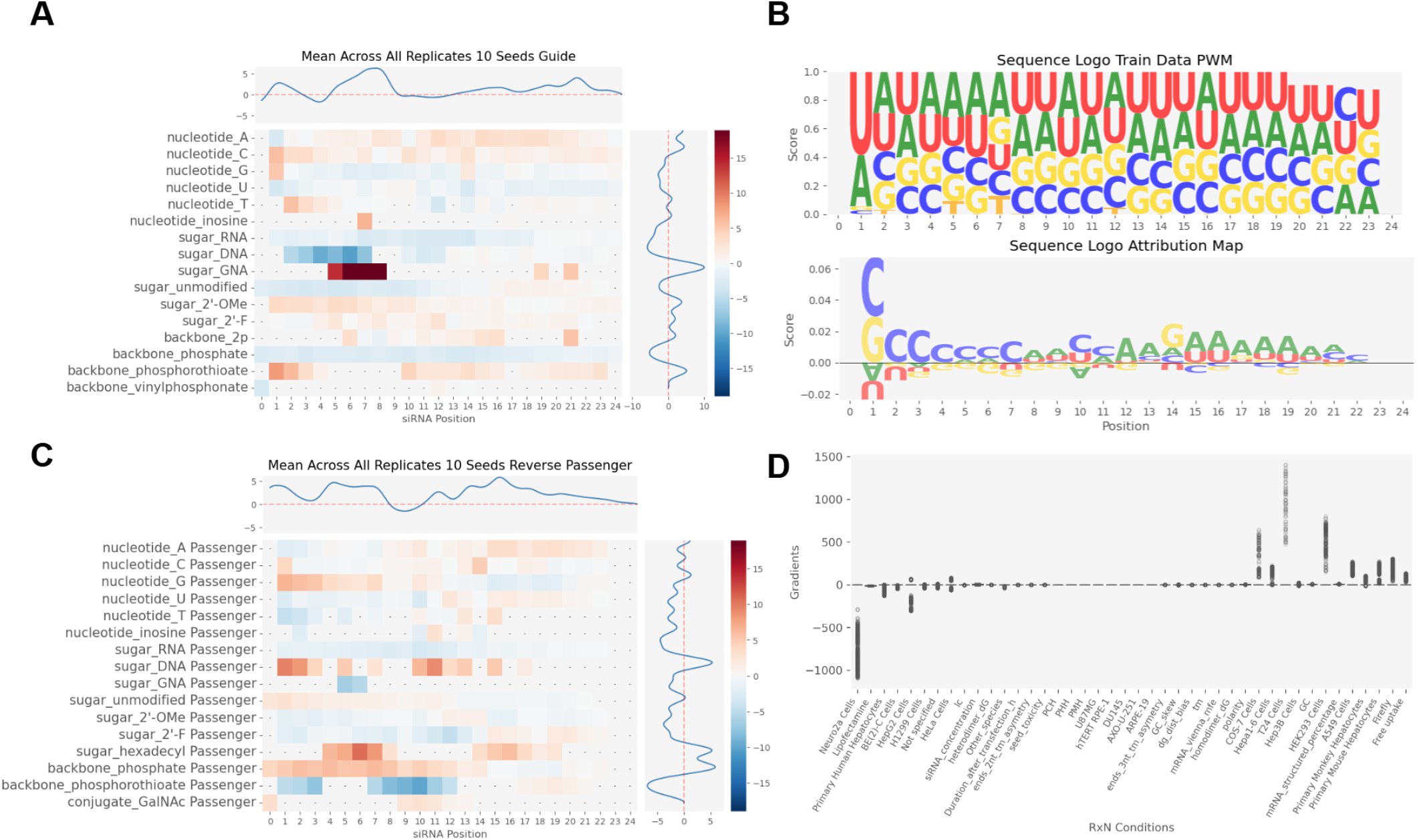
Model interpretation. Integrated gradients attribution maps across sequence and chemistry for (A) guide where red indicates increased predicted mRNA remaining / poor efficacy; blue indicates lower predicted mRNA remaining / strong efficacy. (B) A pairwise weight matrix of observed sequences at particular positions along with corresponding sequence attributions from the sequence where positive numbers are poor efficacy associations, (C) similar to the guide for A but for passengers and (D) meta-feature contributions. All features are normalized by their input frequency x feature mean value

#### Biological interpretation of Orthrus embeddings

To investigate the biological information captured by Orthrus embeddings, we performed principal component analysis (PCA) and correlated the resulting components with interpretable sequence and transcript features (Fig.S2). At the gene level, the first principal component (PC1; 47% of explained variance, Fig. S2C) showed a modest but consistent correlation with RiboNN-predicted translation efficiency across multiple cell types (Spearman *ρ* ≈0.2), while PC2 exhibited a weak association with gene length (Supplementary Fig. S2A). At the target-site level, PC1 correlated more strongly with local mRNA GC content (*ρ* ≈0.4) and inversely with minimum free energy (MFE), indicating that the embedding captures features associated with increased RNA structural stability (Supplementary Fig. S2B). These associations were further supported by lower-variance principal components and mutual information analysis (Supplementary Fig. S3). Overall, the embedding space encodes biologically meaningful properties—including GC content, GC skew, RNA secondary structure, exon junction proximity, and predicted translation efficiency—that are informative beyond gene identity, suggesting that RNA foundation models capture transferable biological signals relevant to siRNA activity.

### In-Silico Design of Modification Patterns

With plausible attribution maps representing known chemical effects, we next asked whether FENNEC could be used as a predictive oracle for siRNA chemical optimization, i.e. to guide the design of more potent siRNAs. To test this, we paired FENNEC with a greedy beam search (Methods) that starts from a known, low-potency parent siRNA and iteratively proposes single-position chemical edits, using FENNEC’s predicted efficacy to select which candidates to keep at each round — a guided “modification walk” through chemical space (Fig. 6A, schematized in Fig. S9) [39]. All six parent sequences used for this in-silico experiment were held out of the standard cross-validation split and had rank-correlation performance above 0.4 Spearman, comparable to the average Spearman of 0.35 across held-out candidates more broadly (Fig. S11) — establishing that FENNEC’s predictions were reasonably well-calibrated for these sequences before using them to guide any search.

**Figure 6.**
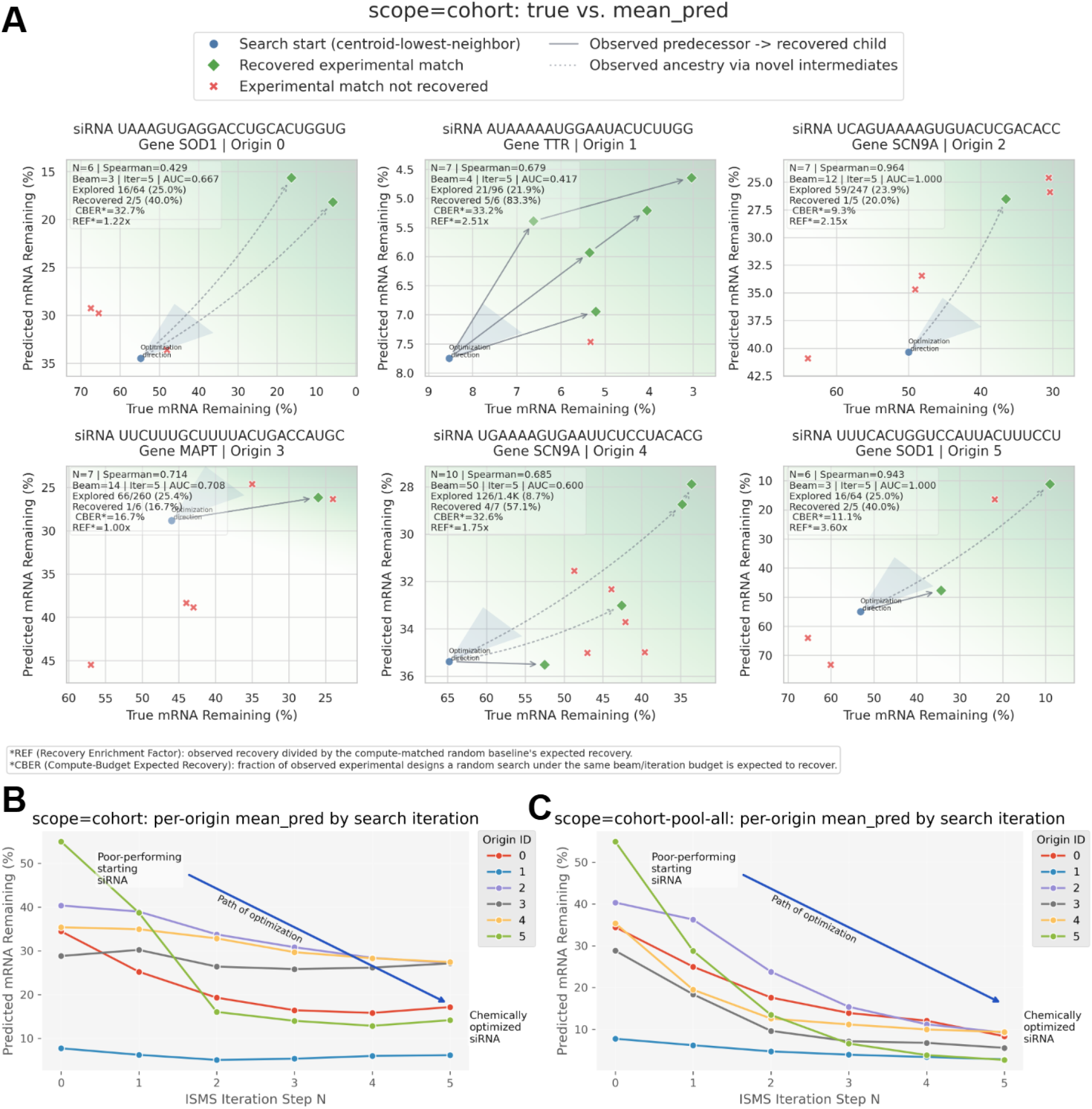
(A) Closed-world (cohort) Ranking Scatterplot of Unique siRNA: Model-predicted efficacy of modification patterns applied to six low-potency parent sequences. The “steepest ascent” trajectory shows the path taken by the greedy beam algorithm, often reaching the upper right of the graph after a few steps; annotations indicate the count of chemically unique patterns and show the recovery of reachable patterns (recovered) within an edit distance of 5 from the origin as well as what a random search would produce, showing the enrichment of using FENNEC as an oracle. (B) Iterative Design (Closed-world cohort): Progression of the optimization walk when restricted to the position-chemistry combinations of exclusively the base siRNA sequences, showing a steady reduction in predicted mRNA %. (C) Open-world (pooled-cohort) Discovery: A broadened scan allowing any modification from any of the 6 base sequence pools to be placed at any position. This illustrates “design headroom,” where the model identifies novel, higher-potency patterns that outperform currently reported chemistry.

We first tested whether this search could rediscover chemistry that already exist in our patent data, using six parent sequences and restricting the search to the pool of positional chemical modifications observed across each base siRNA’s positions (“closed-world” search, see Materials and Methods). Across all six sequences, the search recovered at least one experimentally validated modification pattern (Fig. 6A), despite each search exploring a large space of possible candidates. Notably, FENNEC’s rank correlation with true efficacy within each of these local search spaces was strong, Spearman values ranged from 0.43 to 0.96 across the six sequence families, indicating that FENNEC’s guidance was not just occasionally right but consistently tracked true potency well enough to make the search meaningful. Two examples illustrate the resulting gain over unguided search: for an SCN9A-targeting sequence with 10 known variants (Origin 4, local Spearman = 0.685), FENNEC recovered 4 of 7 reachable variants versus ~2.5 (32.6%) expected by chance (a 1.7-fold enrichment, see Material and Methods); for a SOD1-targeting sequence (Origin 5, local Spearman = 0.943), FENNEC recovered half of the reachable variants versus 16% expected by chance (a 3.1-fold enrichment) — the higher local Spearman here tracking with a cleaner, more reliable search. Beyond simply recovering known chemistry, each round of the beam search progressively increased FENNEC’s predicted efficacy for all six sequences (Fig. ~6B), showing that the search can move toward more potent designs step by step rather than only validating existing ones. Finally, we relaxed the search space to sample from the complete pool of chemical modifications across all six sequence families, rather than restricting each sequence to its own observed chemistry. In this “open-world” setting, FENNEC explored beyond the training distribution to identify novel, high-potency modification patterns for all six sequences. For every sequence tested, the model predicted designs that outperformed all chemistry present in the dataset (Fig. ~6C), highlighting opportunities for siRNA optimization beyond currently reported patent chemistry.

Together, these results indicate that FENNEC can function not only as a predictor but also as a practical design oracle, motivating the experimental validation and design described next.

### Experimental Validation on AHSA1

We assessed the ability of the model to quantitatively predict siRNA-mediated *AHSA1* knockdown across a broad concentration range (0.0006–20 nM; Fig. S7). Predictive performance showed a clear dose dependence. At very low concentrations (≤ 0.006 nM), correlations were weak and not statistically significant, reflecting minimal measurable target suppression at the lowest siRNA dose levels. Within the active concentration range (0.02–6.33 nM), the model demonstrated strong and highly significant predictive power. Spearman correlations ranged from 0.42 to 0.68 (all *P* < 10^−5^), indicating robust recovery of siRNA rank ordering. Linear modeling further showed substantial quantitative accuracy, with predictions explaining up to 43% of the variance in residual mRNA levels at 6.33 nM (*R*^2^ = 0.43, *P* = 6.5 × 10^−13^; Fig. 7A). Performance remained strong at 20 nM (Spearman *r* = 0.62, *R*^2^ = 0.39), with modest attenuation consistent with partial saturation of target knockdown at high concentrations. Across all active concentrations (≥0.02 nM), the mean Spearman correlation was 0.57, indicating stable predictive performance throughout the functional dynamic range of the assay.

**Figure 7.**
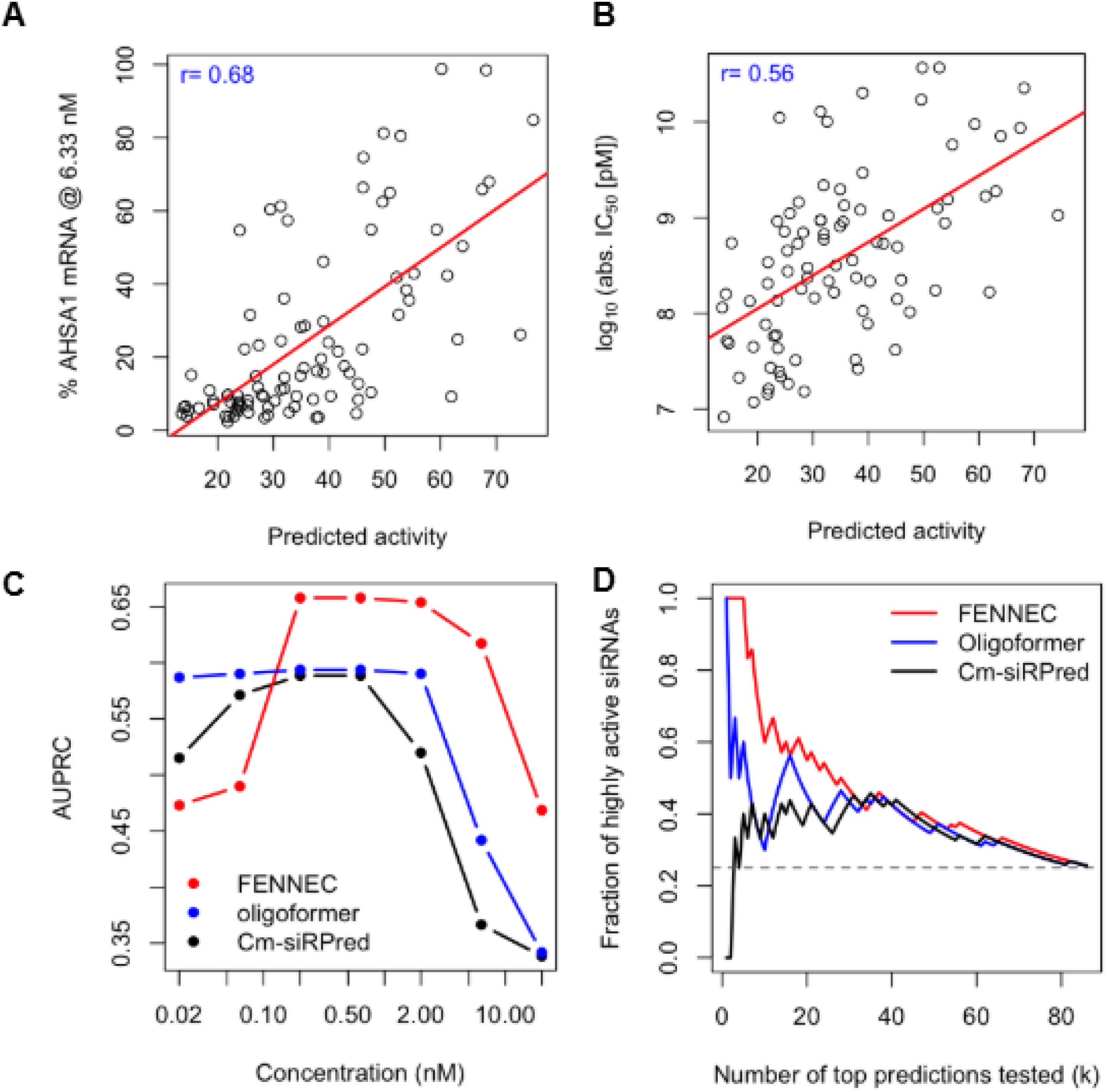
Evaluation of siRNA activity predictions. (A) FENNEC predicted siRNA activity scores plotted against experimentally measured AHSA1 mRNA levels at 6.33 nM. (B) FENNEC predicted activity versus log_10_-transformed absolute IC_50_ values derived from four-parameter logistic fits to dose-response data (*n* = 94). Each point corresponds to an individual siRNA. The red line indicates the least-squares regression fit. (C) Global model performance across siRNA concentrations measured as precision–recall area under the curve (AUPRC). (D) Top-*k* hit rate analysis using residual mRNA levels at 6.33 nM as ground truth. The fraction of highly active siRNAs recovered among the top *k* predicted candidates is shown for each model.

To determine whether the model captures intrinsic siRNA potency rather than concentration-specific readouts, we next correlated predicted residual mRNA with absolute IC_50_ values derived from fitted dose-response curves (Fig. 7B). Predicted mRNA levels showed a strong monotonic association with log-transformed IC_50_ values (Spearman *r* = 0.56), indicating that the model generalizes across the full dose-response profile and captures underlying potency rather than concentration-specific effects.

We next benchmarked FENNEC against Oligoformer and Cm-siRPred using classification-based ranking analyses. Across the tested concentration range, FENNEC consistently achieved the highest precision–recall area under the curve (AUPRC), with peak performance observed at intermediate concentrations (0.2–2 nM; Fig. 7C), where residual mRNA measurements provided the greatest dynamic range for discriminating siRNA activity. For these analyses, highly active siRNAs were defined as the top quartile of sequences with the lowest residual mRNA levels at 6.33 nM, the concentration showing the strongest monotonic relationship between predicted and measured activity. Precision–recall and Top-*k* hit rate analyses at this concentration further showed that FENNEC more effectively prioritized highly active siRNAs among top-ranked predictions than OligoFormer and Cm-siRPred (Fig. 7D and Fig. S6). When potency measurements were used as ground truth, all three models - FENNEC, OligoFormer, and cm-siRNA-pred, enriched potent siRNAs among top-ranked predictions, showing broadly comparable performance across most ranking depths.

## DISCUSSION

### Key Contributions

This work makes four contributions to the field: **(i)** a curated, machine-readable corpus of patent-derived chemically modified siRNAs, spanning an unprecedented diversity of clinically relevant modifications and linked assay metadata across the largest target space analyzed so far in literature; **(ii)** a newly generated data set of 94 chemically modified siRNA for AHSA1 across 10 concentrations for N=940 new data points; **(iii)** a novel, reproducible and interpretable modeling stack for siRNA activity prediction and chemical design with state-of-the-art performance against benchmark models; and **(iv)** evidence that RNA language-model embeddings modestly but consistently improve prediction when combined with explicit chemistry and reaction features. Together, these contributions shorten the loop between design, in-silico screening, and wet-lab experimental validation, which hastens the practical bottleneck in modified-siRNA development.

### Dataset Contributions: Quality and Predictive Value

The availability of large, well-annotated training data has long been identified as the central bottleneck for chemistry-aware siRNA prediction. Here, we address this challenge by developing a rigorous data retrieval and quality control pipeline. Converting patent tables into multimodal sequence/chemistry/assay records is non-trivial at scale: patents vary considerably in table formatting even within a single applicant group, and a substantial fraction of entries could not be reconciled across tables or lacked unambiguous target/transfection context and were excluded. The resulting corpus is, to our knowledge, among the largest compiled sets of chemically modified siRNA with replicate measurements and linked experimental covariates.

The higher replicate counts make the data meaningfully more learnable than prior compilations and expose signals that are attenuated in single-measurement resources. Nevertheless, the curated dataset retains inherent limitations: assay protocols are heterogeneous, concentration series are inconsistent, and certain chemical modifications are disproportionately represented due to the specific focus of the underlying patents. We regard these biases as challenges to be addressed through future data contributions, rather than as fundamental barriers to modeling.

The AHSA1 experimental validation set adds a further 940 datapoints (94 siRNAs across 10 concentrations) generated under controlled, uniform assay conditions, providing a complementary resource with richer concentration coverage than is typical in patent-derived data.

### Relationship to other large scale dataset

One recent resources has sought to address this data gap. While this work was in preparation, the CMsiRNAdb catalog from He et al., [40] was published, a database of 43,153 chemically modified siRNA entries sourced from 90 patents covering 13 therapeutic gene targets and 36 modification types.

CMsiRNAdb substantially increases the volume of publicly available data on chemically modified siRNAs. However, raw entry count is an imperfect proxy for model-relevant data quality, and a direct comparison with our curated dataset reveals a few limitations that constrain its utility for developing broadly generalizable predictive models. For example, as stated above, our curation pipeline specifically enforced quality thresholds that CMsiRNAdb does not: we required ≥95% column datatype purity, a minimum sequence length of 14 nt per strand, and explicit chemical modification exceeding 20% of the sequence.

A more direct comparison reveals that the two resources are largely complementary: only 5 gene targets overlap (APP, CTNNB1, INHBE, MAPT, MARC1) and just 5 of 90 patents are shared. However, CMsiRNAdb explicitly acknowledges a distributional skew in the data: the majority of its entries originate from a small number of cell lines and a narrow set of target genes (PNPLA3 and HSD17B13, primarly metabolic and cardiovascular liver targets), with PNPLA3 comprising 27.5 % of their corpus. The authors themselves caution that “any new predictive models trained directly on this dataset in the future would likely inherit this bias, potentially limiting their generalisation to under-represented cell lines or novel gene targets”. By contrast, the FENNEC patent training set spans 35 patent derived gene targets across diverse therapeutic areas including neurodegeneration (HTT, LRRK2, SOD1), complement biology (C3, CFB, MASP2), coagulation (F10, PLG), pulmonary disease (MUC5B), and rare metabolic disorders (TMPRSS6, HAO1, FLCN), representing a ≈5.8-fold greater target diversity. This represents a more heterogeneous feature landscape from which models can learn transferable signals rather than target-specific features.

CMsiRNAdb contains a greater number of modification types with 36 modification types,including rare chemistry such as locked nucleic acids, 4’-thio sugars, and threofuranosyl nucleotides. However over 85 % of their data employs the same core modification vocabulary (2’-fluoro, 2’-O-methyl, phosphorothioate, GNA, vinylphosphonate, hexadecyl, and GalNAc) captured by our 20 base-agnostic modification classes (combinations of the 12 reported in FENNEC paper), combining mod + backbone/conjugate for comparable reporting across 63 base+mod+backbone/conjugate tokens and 1,134 unique combinatorial patterns. This pattern-level diversity, combined with explicit multi-species annotations (human, cynomolgus macaque, mouse), linked experimental covariates (concentration, cell line, transfection method, duration), and stringent quality filtering, provides the breadth and metadata richness for training generalizable, chemistry-aware predictive models. The two databases are thus best viewed as complementary resources: CMsiRNAdb contributes additional depth for a focused set of targeted genes with exploratory rare chemistry, while the FENNEC corpus provides the target diversity, positional pattern complexity, and experimental context necessary for robust cross-gene, cross-chemistry generalization in siRNA drug design. Both are needed by the community. Future work explicitly reconciling the two, with mutual quality-control pipelines, would be a valuable community contribution.

### Insights from the Experimental Design

Experimental validation on AHSA1 confirms FENNEC’s utility as a design tool in a truly zero-shot regime — a target entirely absent from the training corpus. Across most metrics, chemical modification-aware training provides a clear advantage over competitor models. The one exception is *R*^2^: FENNEC’s predictions span a wider range (e.g. 10–40% mRNA remaining for high-activity siRNAs) than the experimentally observed range (e.g. 5–15%), inflating prediction error even when rank ordering is correct, as reflected by the higher Spearman correlations. FENNEC’s ability to correlate with absolute *IC*_50_ values was an unexpected result, given that no dose-response modelling was included in training, and motivates future incorporation of multi-concentration fitting into the learning objective.

The benchmarking showed measurable differences between the competitor models. Notably, OligoFormer achieved higher *R*^2^ than FENNEC on the AHSA1 validation set (Fig. S6A), reflecting its stronger absolute quantitative calibration in this unmodified-sequence setting. However, OligoFormer lagged behind FENNEC on Spearman rank correlation, a more relevant metric for siRNA prioritisation,indicating that while it captures gross activity levels, it orders sequences less reliably. We attribute OligoFormer’s high performance here compared to the *in silico* benchmarks to (i) the absence of chemical modification diversity in the AHSA1 set, and (ii) the fact that sequences were run with perfect 5’ homology, unlike the JAK1 and APP datasets where 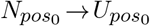 substitution is standard practice. This pattern is consistent with OligoFormer’s architecture: optimised for unmodified sequences, it degrades when the very first guide position deviates from training distribution, as shown in the benchmark analysis.

### Biological Drivers

Interpretability analyses using integrated gradients demonstrate that FENNEC recovers established sequence-level design principles while extending them to the chemically modified siRNA landscape. We mitigate epistemic uncertainty by normalizing and aggregating attributions across multiple seeds. This stabilization reveals plausible, literature-consistent patterns in the saliency maps that might otherwise be obscured by model variance, as well as novel insights.

#### Position Specific Associations

Consistent with established design rules [8, 41, 42], we observe the classic strand asymmetry: AU preference near the 5’ end of the guide (especially position 1), with AU enrichment across the first three positions and a small spike at position 7, and a relatively inverse pattern at the 3’ end. Chemical attributions recapitulate and extend observations reported in the literature [43]. Individual positions show preferences for certain modifications and reveal position-dependent costs — for example, GNA carries a measurable potency penalty, most pronounced at positions 5 and 6 of the guide in our attribution maps — consistent with the known disruptive effect of seed-region GNA on base-pairing and silencing [44]; its use should therefore be deliberate and justified by a specificity constraint rather than applied broadly. Positions 2 and 6 show the most consistent signal: 2’-fluoro modifications are favoured there, aligning with seed-stability tuning and innate-immune quiescence [45]. By contrast, 2’-OMe is bulkier and acts as a steric impediment at positions where the seed engages AGO2, consistent with Foster et al., who attribute potency losses at position 2 to steric clash with the AGO2 MID domain [43]. DNA substitutions are generally tolerated in the 4–7 region but carry a modest negative association at position 14. 2’-fluoro modifications show slightly reduced favourability at positions 5, 11, and 12, possibly reflecting reduced protection against endonucleases at these sites relative to 2’-OMe. More work is needed to fully disentangle these context dependencies, but the attribution maps are consistent with existing literature and provide additional positional nuance for hypothesis generation.

#### Backbone Linkers

Backbone and terminal chemistry behave more uniformly. Phosphorothioates at the terminal positions — particularly near the 3’ end of the passenger, where they enforce strand-loading asymmetry — are consistently beneficial in the derived attributions, matching their known nuclease-protection role. At the guide 5’ end, however, phosphorothioates display a marginally unfavourable effect, consistent with interference with RISC loading. Research from Alnylam and others [46, 47] attributes this to the racemic mixture of *R*_*p*_ and *S*_*p*_ diastereomers produced during synthesis, with the *S*_*p*_ form hindering RISC loading; selective exclusion of the *S*_*p*_ diastereomer can actually increase knockdown.

#### Reaction Conditions

Reaction and meta-features contribute measurable, albeit modest, predictive value, likely functioning in part as batch-correction variables that account for systematic differences between assay platforms. Cell line, transfection method, duration, and log-transformed siRNA concentration all contribute predictive signal, while engineered thermodynamic features contribute more selectively across regimes. The practical implication is clear: models deployed to new assay systems benefit from incorporating these covariates and, ideally, from fine-tuning on a small number of system-specific examples (e.g., 10–50) to ensure accurate calibration.

#### Foundation Models and Transfer Learning

The benefit of FM embeddings was most pronounced where target-sequence context was genuinely novel. ACVR1C, with an atypical 3’ UTR architecture poorly represented in training, showed near-chance zero-shot performance without FM embeddings; incorporating either Orthrus or RNA-FM recovered predictive signal with markedly fewer labeled examples than the FM-ablated model required. More broadly, FENNEC demonstrated strong transfer to novel chemistry even without FM support: JAK1, featuring an out-of-distribution alternating 2’-OMe/2’-F pattern absent from training, achieved reasonable zero-shot accuracy and reached high correlation with modest fine-tuning. Crucially, FM embeddings provided no meaningful benefit for JAK1, in stark contrast to ACVR1C. This dissociation supports a practical design principle: FM embeddings are most valuable when the prediction bottleneck is target-sequence novelty, while the explicit chemistry encoding is sufficient when the bottleneck is an unfamiliar modification pattern.

A natural concern is that Orthrus, trained on mRNA sequences, could improve predictions simply by encoding a high-dimensional proxy for gene identity, in effect, memorising which target is being silenced rather than capturing any transferable sequence property. Although target identity is an important factor in siRNA design and is substantially represented within the Orthrus embedding (Fig. S4), several observations suggest that the embedding captures biological signals beyond gene identity alone. First, even when restricting to the siRNA-centred 100 bp slice alone (excluding UTR regions), Orthrus still outperforms the FM-ablated model, suggesting the benefit is not purely driven by gene-level context. Second, because RNA-FM operates without target-level supervision yet confers similar data-efficiency gains, the advantage is attributable to shared low-level sequence and structural features rather than target memorisation. Third, principal-component analysis of the embedding space (Supplementary Fig. S2–S4) shows that leading dimensions correlate with interpretable biophysical properties — including GC content, GC skew, Vienna-calculated MFE of the siRNA-centred mRNA window, exon junction proximity, and RiboNN-predicted translation efficiency across tissues — all of which are informative independently of gene identity. Together, these results suggest that RNA foundation models contribute genuine, transferable biological signal and motivate their inclusion particularly when predicting activity on novel gene targets or non-standard chemical scaffolds.

### In silico Guided Design

Our in silico design experiments, run on sequences spanning a range of predicted potency, support two main conclusions. First, FENNEC can rank siRNAs by predicted efficacy from chemical modification patterns alone. Second, it can actively guide sequence diversification: a greedy beam search steered by FENNEC recovered chemically valid, patent-derived modification patterns far more efficiently than random exploration, with a Recovery Enrichment Factor ranging from near 1 (no advantage, for sequences with few tested variants) to 3.86x for a TTR-targeting sequence. This demonstrates that model guidance is meaningful even with simple generative algorithms. This claim is in silico and not yet backed by a prospective wet-lab experiment; we regard it as a validated design hypothesis rather than a confirmed design tool. Fig.6B– C further show that expanding to an open-world modification pool allows the search to reach a lower predicted-efficacy minimum more rapidly at the same beam width and iteration count.

### Benchmarking against the State-of-the-Art

Our benchmarking results underscore two critical findings for the field. First, the superior performance of deep learning architectures over ensemble tree methods (XGBoost) suggests that the mapping from chemical modification patterns to biological efficacy involves high-order conceptual interactions that simpler models capture less effectively as in the case of the linear regressors, despite good baseline performance. Second, the comparison with existing tools reveals a significant gap in current capabilities: while sequence-only models like RNAxs and Oligoformer offer valuable baselines, their difficulty to natively process diverse chemical modifications (or even standard industrial scaffolds like 5’-U replacement) limits their utility in modern drug design. FENNEC’s ability to outperform even chemistry-aware models like cm-siRNA-pred highlights the importance of training data scale and quality, which we believe is the primary factor underlying FENNEC’s advantage over comparable architectures. By explicitly encoding chemical identity and position alongside foundation model embeddings, FENNEC effectively bridges the gap between traditional sequence-based rules and the complex reality of chemically modified therapeutics.

## Limitations

While FENNEC advances the state of the art, it faces three key limitations. First (i), the training data, derived primarily from patents, is inherently biased toward successful chemical scaffolds, potentially limiting the model’s ability to identify novel, structurally distinct modification patterns. Second (ii), our use of one-hot encoding, despite being effective for representing chemical modifications, treats distinct chemistry (e.g., 2’-F vs. 2’-OMe) as categorical labels; this approach could be missing some nuance to explicitly capture stereochemical or biophysical properties (e.g., hydrophobicity, solvent accessibility) that could have an enhanced representation via molecular graph embeddings (GNNs). Finally (iii), there remains a translational gap between the *in vitro* data used for training and true *in vivo* therapeutic efficacy, which involves complex pharmacokinetic barriers not modeled here. Methodologically, our encodings capture modification identity and position, but not full 3-D or solvent context; moving to richer representations presents a promising avenue for improvement. Finally, acknowledging that integrated gradients are sensitive to baseline selection, we validated our findings using finite-difference perturbations and recommend this dual-verification approach for downstream applications.

## Conclusion

In summary, we provide an experimentally validated foundation for chemistry-aware siRNA design: a curated patent-derived training corpus, a fast and competitive architecture that leverages RNA LLM embeddings, and interpretable maps that both reproduce known sequence rules and highlight nuanced, context-dependent chemistry effects. The most actionable guidance is to treat the modifications as key design points; to make passenger-strand and terminal chemistry important considerations; and to allocate a small budget for system-specific calibration.

## ACKNOWLEDGMENTS

We would like to thank Johannes Linder of Helmholtz-Munich for their contributions on project scope and scientific conceptualization.

## Author contributions

Data curation, analysis, validation, and computational experiments, A.L; Compound synthesis and experimental validation, J.B., P.B., D.Y; Conceptualization, funding acquisition, project administration, and supervision, A.M. and D.B.; Contribution to method conceptualizaiton and interpretation of results, R.R., J.B.; Writing original draft, A.L., A.M; Review and editing, all authors.

## DATA AVAILABILITY

Source patents referenced in this study are publicly available through Zenodo at https://zenodo.org/records/21824464 with DOI:10.5281/zenodo.21824464.

## CODE AVAILABILITY

The FENNEC model architecture, inference scripts, and associated processing utilities are available on GitHub at: https://github.com/marsico-lab/fennec.

