## Supplementary for "FENNEC: Fine-Tuned Ensemble Neural Networks Accelerate Chemically Modified siRNA Screening and Design"

### SUPPLEMENTARY MATERIAL

**Table S1.** Formulas for `additional engineered features on` $s-(Seq_{unmodified})$ and $m-(mRNA))$

| Feature | Formula / Symbol | Meaning | Units / Range |
|---|---|---|---|
| homodimer_dG | $\Delta G_{\text{homo}}(s)$ | Primer3 homodimer $\Delta G$ at standard ionic conditions | kcal/mol |
| heterodimer_dG | $\Delta G_{\text{hetero}}(s,m)$ | Primer3 heterodimer $\Delta G$ between $s$ and $m$ | kcal/mol |
| seed_toxicity | $\text{Tox}\big(s_{\text{U}}[2{:}7]\big)$ | Lookup in seed-toxicity table (positions 2–7; $T\to U$) | dataset-defined |
| tm | $\text{Tm}(s)$ | Nearest-neighbor melting temperature at standard conditions | °C |
| GC | $\dfrac{\#G+\#C}{L}$ | GC fraction of $s$ | 0–1 |
| ends_3nt_tm_asymmetry | $\text{Tm}(s_{1..3})-\text{Tm}(s_{L-2..L})$ | $5'$ vs. $3'$ Tm difference (3-nt windows) | °C |
| ends_2nt_tm_asymmetry | $\text{Tm}(s_{1..2})-\text{Tm}(s_{L-1..L})$ | $5'$ vs. $3'$ Tm difference (2-nt windows) | °C |
| dg_dist_bias | $\sum_{i=0}^{L-2}\Delta G(r_i r_{i+1})w(i,L)$ | Weighted $\Delta G$ nearest-neighbor sum after $T\to U$; $w(i,L)=\alpha d^2\dfrac{d}{\lvert d\rvert+1}$ with $\alpha=\dfrac{3.5}{(\frac{L}{2}+1)^2}$, $d=\frac{L}{2}+1-i$ | kcal/mol |
| polarity | $\frac{1}{L}\sum_{i=1}^{L}B(s_i)$ | Mean B-factor per base (Kirillova 2011 mapping) | a.u. |
| GC_skew | $\dfrac{\#G-\#C}{\#G+\#C}$ | GC skew of $s$ - sequence | $[-1,1]$ |
| mRNA_structured_percentage | $\text{unp}(m)$ | Fraction of unpaired nucleotides from ViennaRNA fold | fraction or % |
| mRNA_vienna_mfe | $\text{MFE}(m)$ | Minimum free energy from ViennaRNA fold | kcal/mol |
| linguistic_complexity | $\dfrac{U_k(s)}{\min(4^k,L-k+1)}$, $k=4$ | Linguistic complexity over $k$-mers; $U_k(s)$ = number of distinct $k$-mers in $s$ | 0–1 |

**Table S2.** Concentration-dependent correlation between predicted siRNA activity and measured AHSA1 knockdown. Spearman correlation coefficients and linear regression statistics between predicted activity scores (FENNEC) and experimentally measured residual AHSA1 mRNA levels across the tested siRNA concentration range. $n$ denotes the number of siRNAs analyzed at each concentration.

| concentration [nM] | $n$ | spearman_r | spearman_p | $R^2$ | slope | lm_p |
|---:|---:|---:|---:|---:|---:|---:|
| 0.000637 | 94 | 0.18 | 0.075925 | 0.02 | 0.11 | 0.236028 |
| 0.00201 | 94 | 0.13 | 0.207045 | 0.03 | 0.14 | 0.091284 |
| 0.00636 | 94 | 0.18 | 0.086372 | 0.02 | 0.11 | 0.210486 |
| 0.0201 | 94 | 0.42 | 3.16E-05 | 0.16 | 0.58 | 6.02E-05 |
| 0.0635 | 94 | 0.44 | 9.72E-06 | 0.19 | 0.79 | 1.49E-05 |
| 0.201 | 94 | 0.59 | 0 | 0.32 | 1.24 | 2.35E-09 |
| 0.634 | 94 | 0.59 | 3.35E-10 | 0.31 | 1.17 | 5.84E-09 |
| 2 | 94 | 0.65 | 0 | 0.34 | 1.12 | 6.43E-10 |
| 6.33 | 94 | 0.68 | 0 | 0.43 | 1.06 | 6.54E-13 |
| 20 | 94 | 0.62 | 0 | 0.39 | 0.88 | 1.23E-11 |

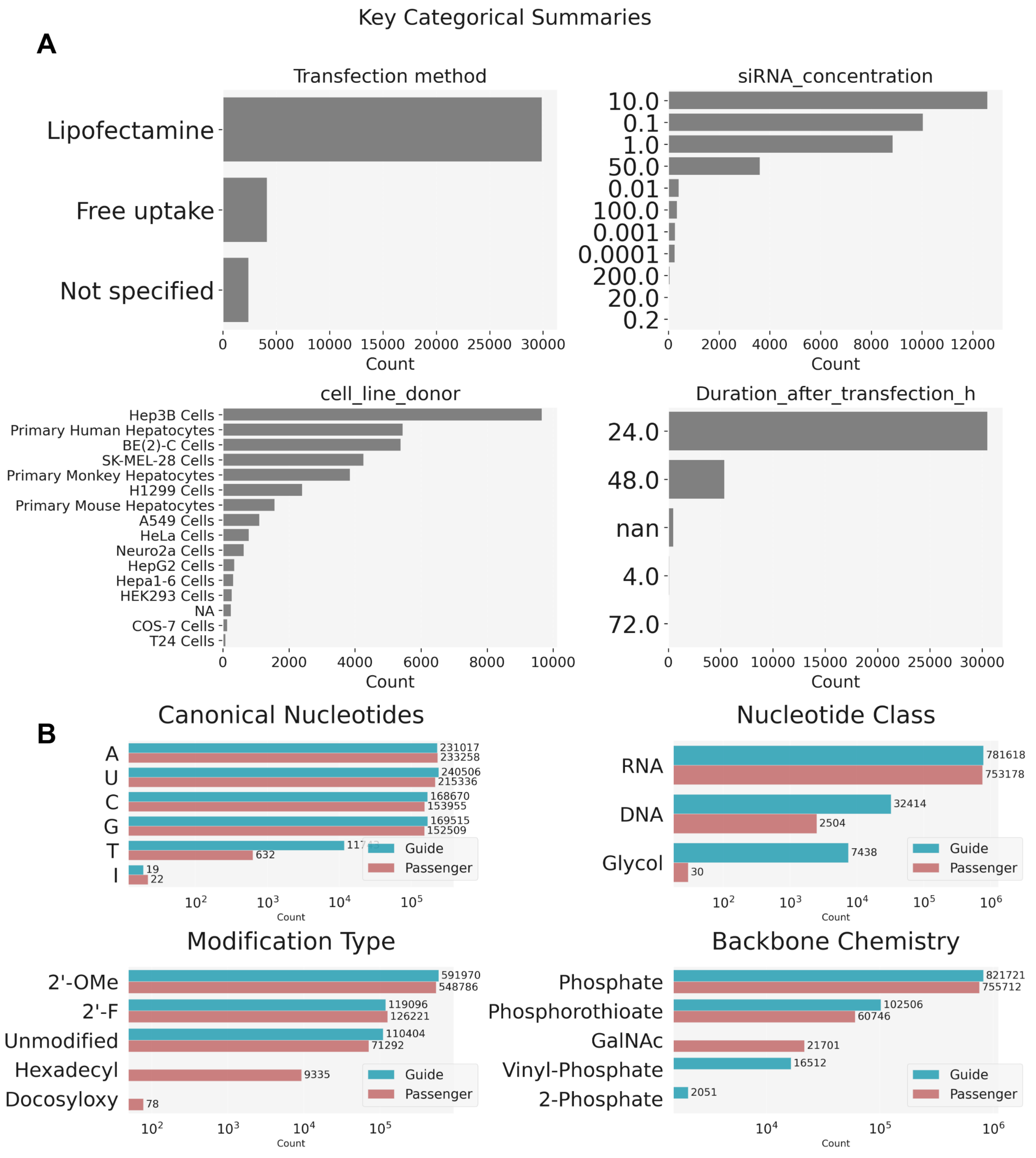


**Figure S1.** Extended Sequence and Experimental conditions graphs (A) Assay/meta-context composition: transfection method, siRNA concentration (binned), cell line donor species, and evaluation time post-transfection. (B) which chemically modified nucleotides are represented most in the data. Covariate heterogeneity motivates inclusion of reaction/meta feature tracks. S1

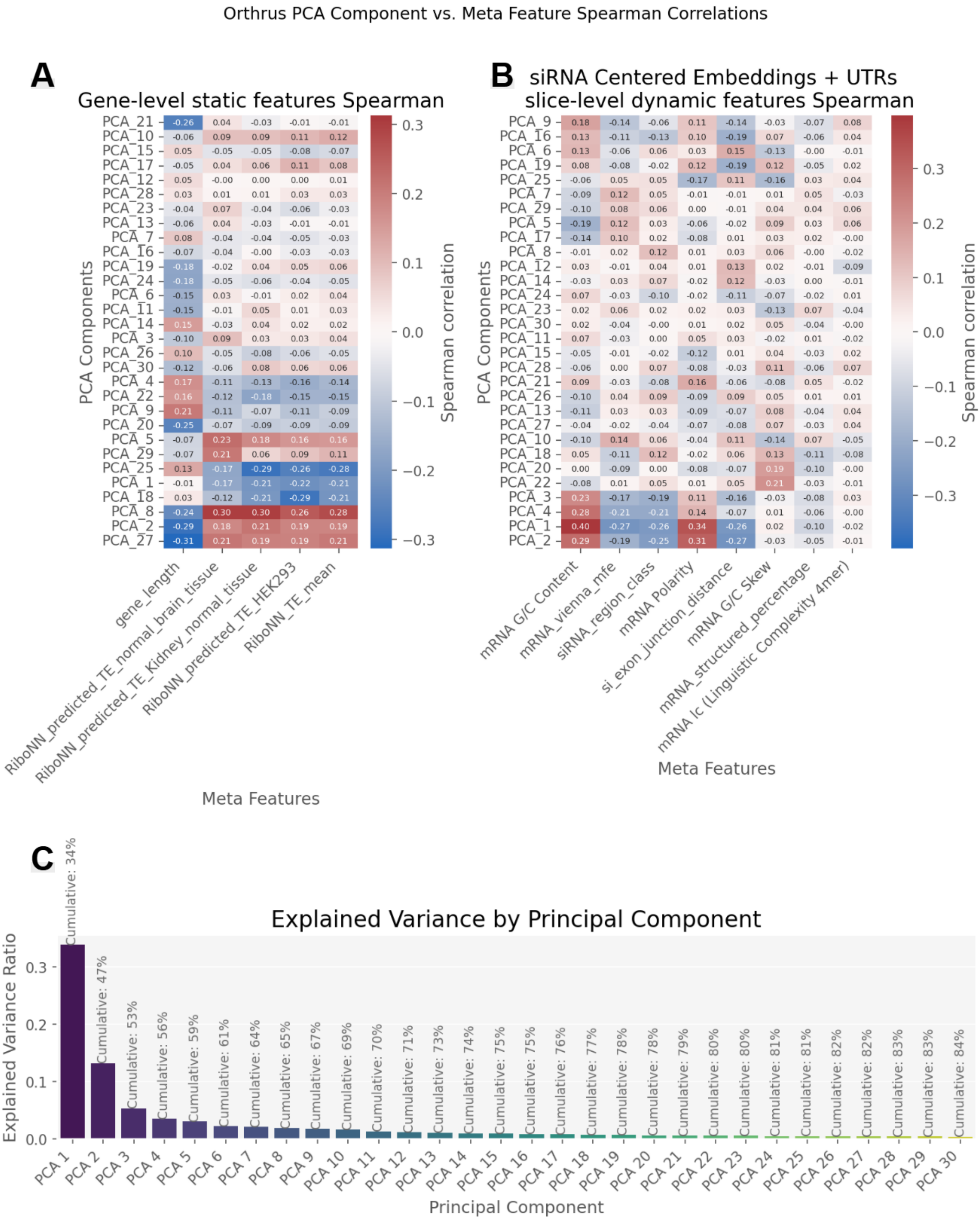


**Figure S2.** Orthrus Embedding Correlations. Analysis of Orthrus embedding dimensions and their correlation/attribution to engineered features, Including gene length, RiboNN predicted TE, and exon distance. (A) shows the correlation of the median principal component given gene X, as these calculated values do not change per gene. (B) correlates the PCA embeddings for every usage 5'UTR 100bp + siRNA-centered mRNA slice 100bp + 3' UTR beginning 100bp slice, All features in B are calculated on the siRNA centered mRNA 100bp slice. (C) The explained variance across the PCA components, totaling just under 85% by component 30. Components used in C are the same used for S3

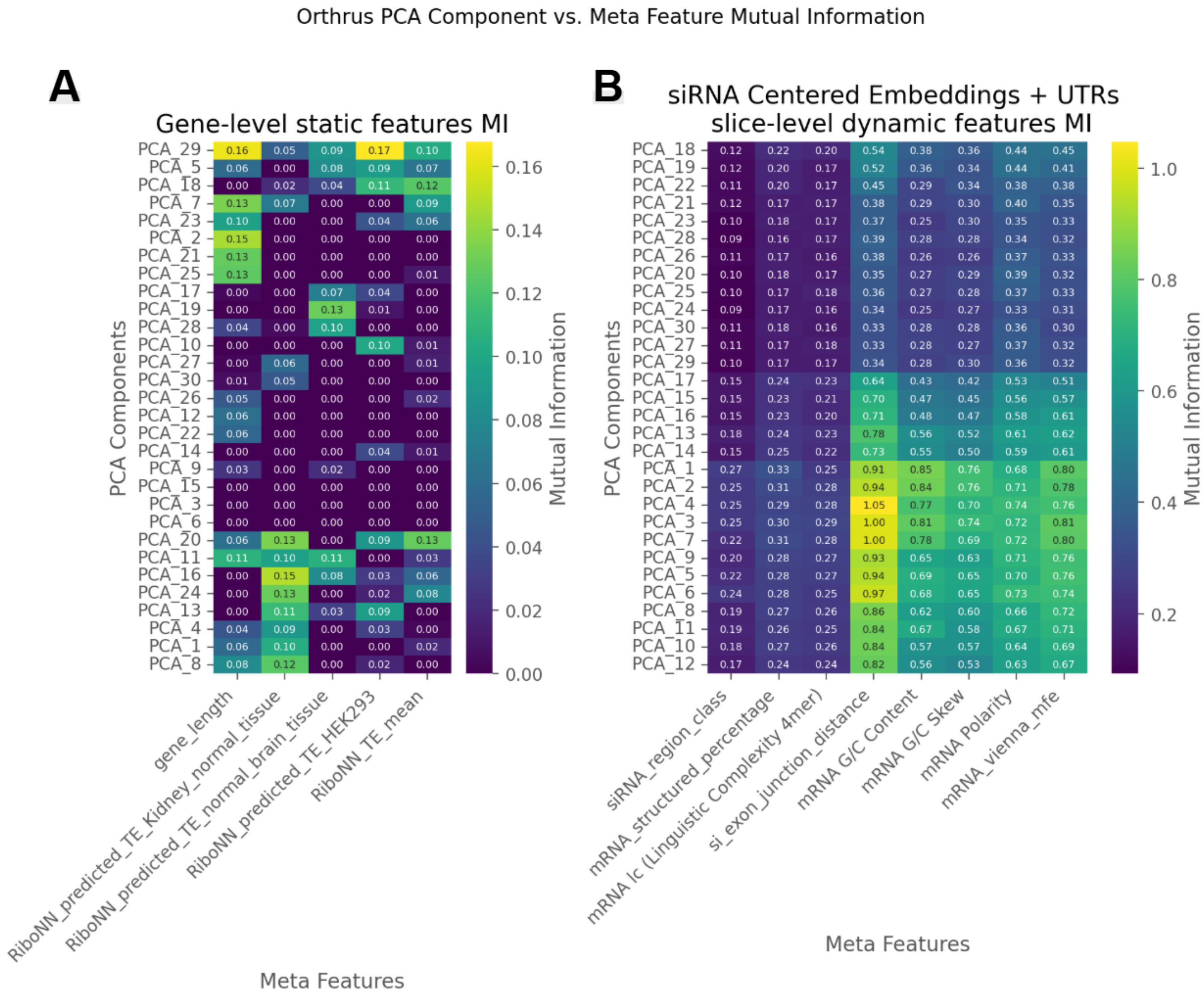


**Figure S3.** Orthrus Embedding Mutual information. Analysis of Orthrus embedding dimensions and their shared mutual information, Including gene length, RiboNN predicted TE, and exon distance. (A) shows the correlation of the median principal component given gene X, as these calculated values do not change per gene. (B) correlates the PCA embeddings for every usage 5’UTR 100bp + siRNA-centered mRNA slice 100bp + 3’ UTR beginning 100bp slice, All features in B are calculated on the siRNA centered mRNA 100bp slice.

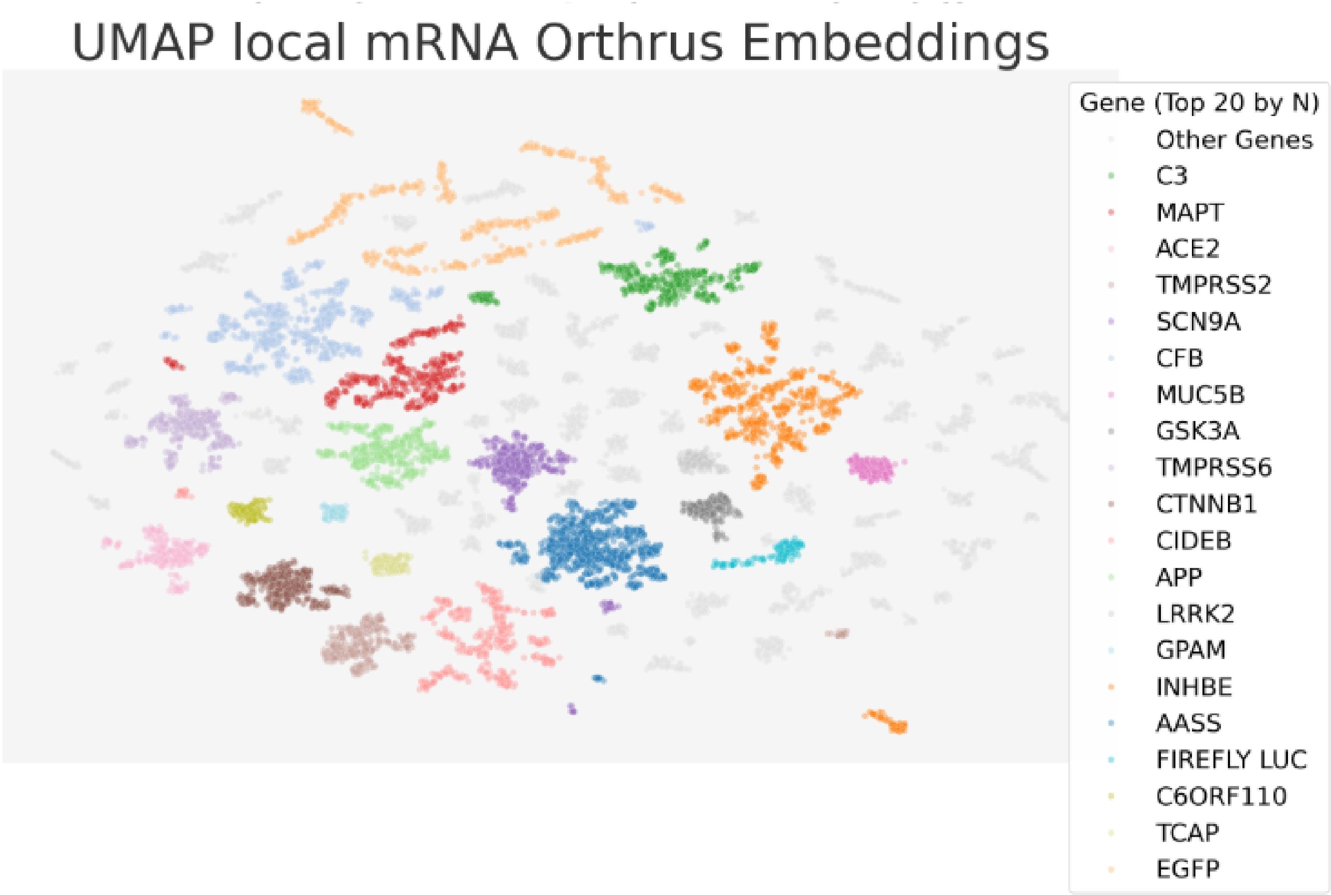


**Figure S4.** Principal Component Analysis (PCA) of Orthrus Embeddings. Visualizing the latent space structure captured by the foundation model colored by gene transcript.

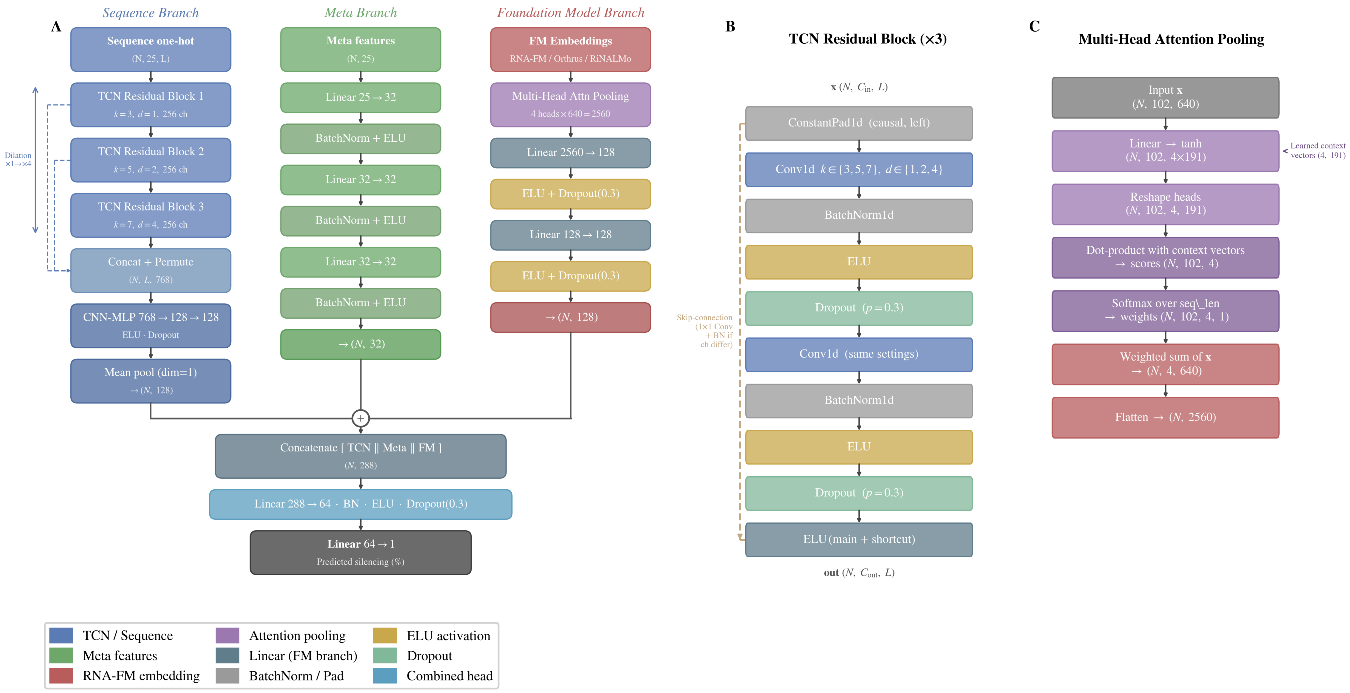


**Figure S5.** Detailed Model Architecture. Extended diagram showing layer-specific details of the FENNEC framework.

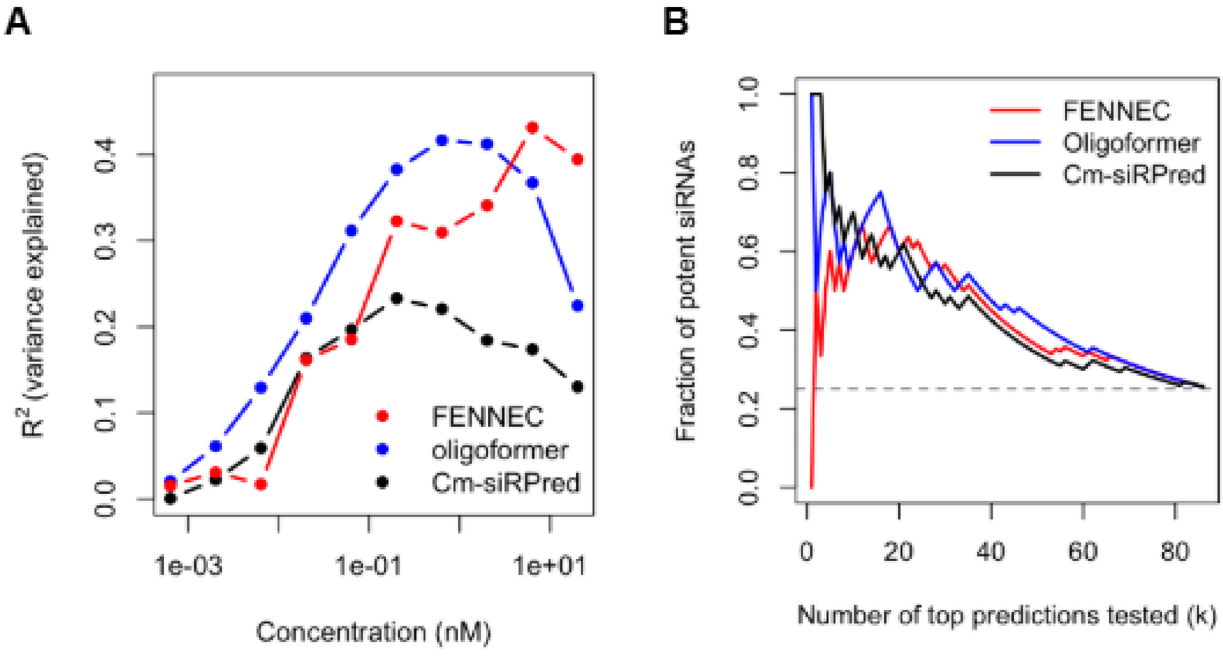


**Figure S6.** Benchmarking of siRNA activity prediction models. (A) Predictive performance of different siRNA activity predictors across concentrations (0.0006–20 nM), quantified as the proportion of explained variance ($R^2$) from linear regression of measured knockdown on predicted activity. (B) Top-$k$ hit rate analysis using absolute IC50 derived from dose-response curves.

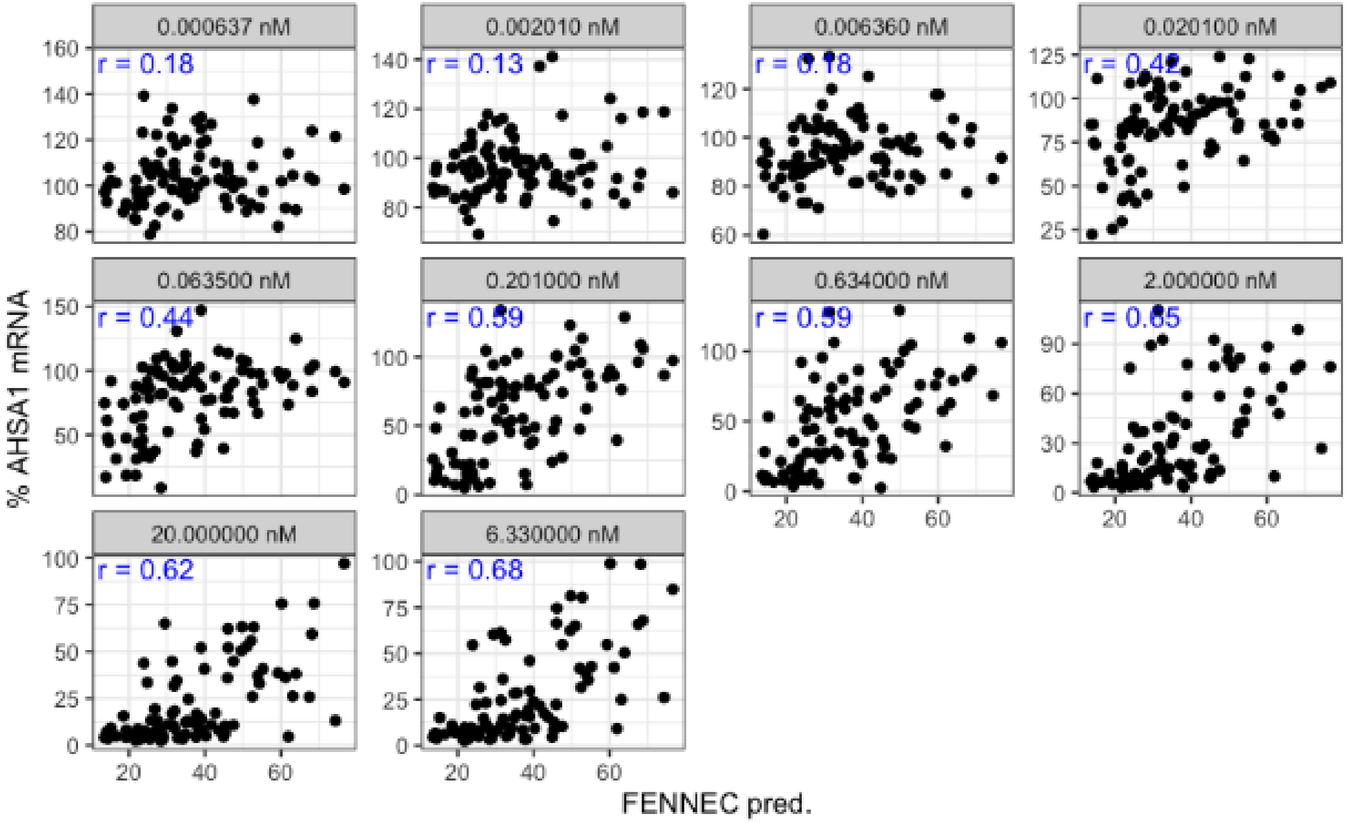


**Figure S7.** Concentration-dependent correlation between FENNEC predicted siRNA activity and experimental AHSA1 knockdown. Predicted siRNA activity were plotted against the percentage of AHSA1 mRNA remaining at the indicated concentrations. Each point represents one siRNA. Spearman rank correlation coefficients ($r$) are indicated in each panel. Lower percentages of mRNA remaining correspond to greater target knockdown.

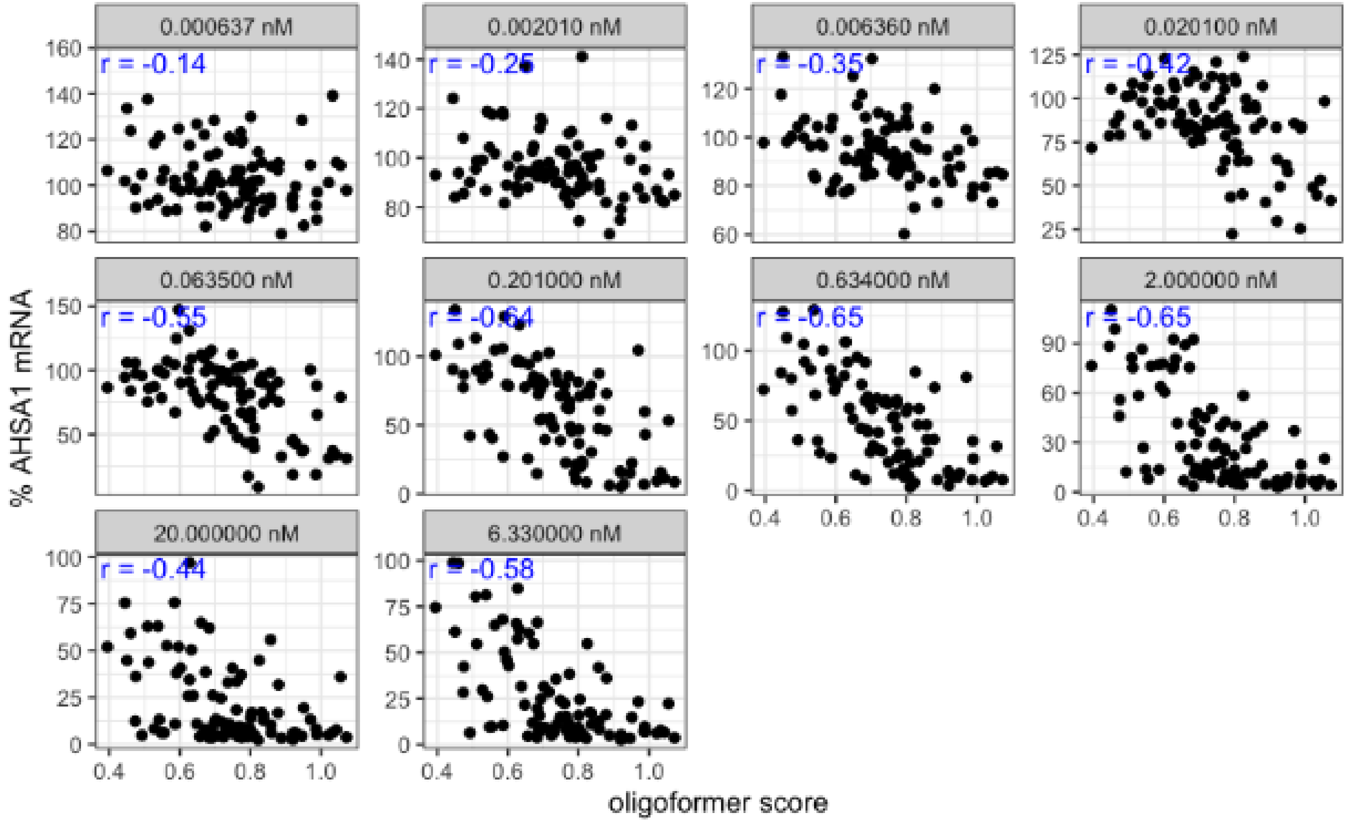


**Figure S8.** Concentration-dependent correlation between Oligoformer activity score and experimental AHSA1 knockdown. siRNA activity scores were plotted against the percentage of AHSA1 mRNA remaining at the indicated concentrations. Each point represents one siRNA. Spearman rank correlation coefficients ($r$) are indicated in each panel. Lower percentages of mRNA remaining correspond to greater target knockdown.

Table S3: Summary of Datapoint Counts per Patent Table (Landscape, 50 chars)

| Patent ID | Table Title | meta_group | Gene | Data (N) |
|---|---|---|---|---|
| WO-2025064660-A2 | Table 6. Single dose screen for dsRNA agents targe... | Patent | ACVR1C | 4179 |
| WO-2023076451-A1 | Table 4. In Vitro Single dose Screens in Primary H... | Patent | CFB | 1982 |
| WO-2025034560-A1 | TABLE 7. Multi-dose In vitro Screening of AASS dsR... | Patent | AASS | 1966 |
| WO-2023003922-A1 | Table 21A. Single dose screen for dsRNA agents tar... | Patent | INHBE | 1939 |
| WO-2025034560-A1 | TABLE 9. Multi-Dose In vitro Screen Free-Uptake of... | Patent | AASS | 1443 |
| WO-2021207189-A1 | Table 8. SCN9A in vitro multidose-dose screen with... | Patent | SCN9A | 1365 |
| WO-2023034837-A2 | Table 7. CIDEB Single Dose Screens in Hep3B Cells | Patent | CIDEB | 1273 |
| WO-2020132227-A2 | Table 4. APP Single Dose Screen in Primary Cynomol... | Patent | APP | 1069 |
| WO-2021207189-A1 | Table 17. SCN9A in vitro multidose-dose screen wit... | Patent | SCN9A | 989 |
| WO-2023003995-A1 | Table 4. Single Dose Screens in Hep3b Cells | Patent | CTNNB1 | 954 |
| WO-2022231999-A1 | Table 8. Single Dose Screen in Hep3b Cells | Patent | TMPRSS6 | 881 |
| WO-2021206922-A1 | Table 4. TMPRSS2 Single Dose Screens in PHH cells | Patent | TMPRSS2 | 874 |
| WO-2025034560-A1 | TABLE 8. Multi-Dose In vitro Screen Free-Uptake of... | Patent | AASS | 862 |
| WO-2021206917-A1 | Table 6. ACE2 Single Dose Screens in PHH cells | Patent | ACE2 | 797 |
| Not From Patent | No Table Title | Takayuki | EGFP | 702 |
| WO-2021202511-A2 | Table 24. MAPT Single Dose Screens in BE(2)C Cells... | Patent | MAPT | 588 |
| WO-2023278607-A1 | Table 8. Dose Screens of LRRK2 dsRNA Agents (L96-C... | Patent | LRRK2 | 455 |
| WO-2021081026-A1 | Table 33. C3 Transfection Single Dose Screens in P... | Patent | C3 | 429 |
| WO-2023049871-A2 | Table 7. In vitro screen of human MAPT siRNA | Patent | MAPT | 397 |
| WO-2020132227-A2 | Table 7. APP Single Dose Screen in Primary Mouse H... | Patent | APP | 356 |
| WO-2022174000-A2 | Table 21. Superoxide Dismutase 1 In Vitro Single D... | Patent | SOD1 | 343 |
| WO-2023014677-A1 | Table 7. Single Dose In Vitro Screens in Hep3B Cel... | Patent | TTR | 323 |
| WO-2024077148-A1 | Table 11. Initial in vitro screening assay in PHH ... | Patent | PLG | 294 |
| WO-2021081026-A1 | Table 12. C3 Single Dose Screens in Hep3B cells | Patent | C3 | 292 |
| WO-2020132227-A2 | Table 18. APP Dose Screen Study in Neuro2A Cell Li... | Patent | APP | 269 |
| WO-2021252649-A2 | Table 7. In vitro screen of human GPAM siRNA in He... | Patent | GPAM | 268 |
| WO-2022174000-A2 | Table 14. Superoxide Dismutase 1 In Vitro Single D... | Patent | SOD1 | 260 |
| WO-2021202511-A2 | Table 10. MAPT Single Dose Screens in BE(2)C Cells... | Patent | MAPT | 249 |
| WO-2022087329-A1 | Table 8. MUC5B Single Dose In Vitro Screen in A549... | Patent | MUC5B | 249 |
| WO-2023014677-A1 | Table 4. Single Dose In Vitro Screens in Primary C... | Patent | TTR | 248 |
| WO-2022174000-A2 | Table 22. Superoxide Dismutase 1 In Vitro Single D... | Patent | SOD1 | 248 |
| WO-2021222549-A1 | Table 8. Complement Factor B In Vitro Single Dose ... | Patent | CFB | 245 |
| WO-2020132227-A2 | Table 17. APP Dose Screen Study in Be(2)C Cell Lin... | Patent | APP | 238 |
| WO-2023044370-A2 | Table 6. In Vitro C3 Single Dose Screen in Hep3b C... | Patent | C3 | 226 |
| WO-2023003995-A1 | Table 7. Single Dose Screens in Hep3b Cells | Patent | CTNNB1 | 225 |
| WO-2021081026-A1 | Table 26. C3 Single Dose Screens in PCH cells (% C... | Patent | C3 | 224 |
| WO-2021081026-A1 | Table 13. C3 Single Dose Screens in PMH cells | Patent | C3 | 224 |
| WO-2021081026-A1 | Table 32. C3 Free Uptake Single Dose Screens in PC... | Patent | C3 | 219 |
| WO-2021222549-A1 | Table 11. Complement Factor B In Vitro Single Dose... | Patent | CFB | 123 |
| WO-2022232343-A1 | Table 4. In Vitro STAT6 Single Dose Screen in A549... | Patent | STAT6 | 122 |
| WO-2023034837-A2 | Table 8. CIDEB Single Dose Screens in Hep3B Cells | Patent | CIDEB | 118 |

Continued on next page

Table S3: Summary of Datapoint Counts per Patent Table (Landscape, 50 chars)

| Patent ID | Table Title | meta_group | Gene | Data (N) |
|---|---|---|---|---|
| WO-2014190157-A1 | Table 6. TMPRSS6 single dose screen. | Patent | TMPRSS6 | 117 |
| WO-2022174000-A2 | Table 15. Superoxide Dismutase 1 In Vitro Single D... | Patent | SOD1 | 117 |
| Not From Patent | No Table Title | Huesken | MMP7 | 113 |
| WO-2022076291-A1 | Table 4. In Vitro Single Dose Screen in Hepa1-6 Ce... | Patent | GPR75 | 106 |
| WO-2014190157-A1 | Table 9. TMPRSS6 Single Dose Screen | Patent | TMPRSS6 | 104 |
| WO-2021081026-A1 | Table 10. C3 Single Dose Screens in Hep3B cells | Patent | C3 | 96 |
| WO-2021087036-A1 | Table 34. HTT Single Dose Screens in BE(2)C Cells | Patent | HTT | 96 |
| WO-2021081026-A1 | Table 11. C3 Single Dose Screens in PMH cells | Patent | C3 | 95 |
| WO-2021222549-A1 | Table 12. Complement Factor B In Vitro Single Dose... | Patent | CFB | 93 |
| Not From Patent | No Table Title | Huesken | P2RX3 | 90 |
| Not From Patent | No Table Title | ichihara_mix | CYLOPHILIN B | 90 |
| WO-2015089368-A2 | Table 10. CFB single dose screen in Primary Mouse ... | Patent | CFB | 88 |
| WO-2015089368-A2 | Table 9. CFB single dose screen in Hep3B Cells | Patent | CFB | 88 |
| WO-2021150969-A1 | Table 5. LRRK2 Single Dose Screens in A549 Cells | Patent | LRRK2 | 79 |
| WO-2016057893-A1 | Table 3b. Additional HAO1 single dose screen in pr... | Patent | HAO1 | 78 |
| WO-2025064660-A2 | Table 13. Single Dose Screen of dsRNA Agents Targe... | Patent | ACVR1C | 78 |
| WO-2015089368-A2 | Table 15. C3 Single dose screen in Hep 3B cells | Patent | C3 | 78 |
| WO-2021087036-A1 | Table 16. HTT Single Dose Screens in BE(2)C Cells | Patent | HTT | 75 |
| WO-2015089368-A2 | Table 14. C3 Single dose screen in Primary Mouse H... | Patent | C3 | 75 |
| WO-2022178411-A1 | Table 12. In vitro screen of mouse FLCN siRNA in P... | Patent | FLCN | 72 |
| WO-2021202511-A2 | Table 9. MAPT Single Dose Screens in BE(2)C Cells-... | Patent | MAPT | 71 |
| WO-2021087036-A1 | Table 4. HTT Single Dose Screens in BE(2)C Cells | Patent | HTT | 69 |
| WO-2021081026-A1 | Table 27. C3 Single Dose Screens in PCH cells (% C... | Patent | C3 | 67 |
| WO-2019014530-A1 | TABLE 6A. Single dose screen in Primary Mouse Hepa... | Patent | LDHA | 66 |
| WO-2022087329-A1 | Table 11. MUC5B Single Dose In Vitro Screen in Cos... | Patent | MUC5B | 63 |
| WO-2014190157-A1 | Table 11. TMPRSS6 Single Dose Screen | Patent | TMPRSS6 | 61 |
| WO-2023056478-A1 | Table 8. Mouse ANGPTL7 siRNA dose screen in COS-7 ... | Patent | ANGPTL7 | 58 |
| WO-2022178411-A1 | Table 13. In vitro screen of human FLCN siRNA in A... | Patent | FLCN | 58 |
| WO-2021257568-A1 | Table 5. ALK Single Dose Screen in Hepa1-6 Cells | Patent | ALK | 51 |
| WO-2024077148-A1 | TABLE 10. PLG Multi-Dose Screens in Primary Cyno H... | Patent | PLG | 50 |
| WO-2024077148-A1 | TABLE 9. PLG Multi-Dose Screens in Primary Human H... | Patent | PLG | 50 |
| WO-2021237097-A1 | Table 6. MARC1 in vitro multi-dose screen with a s... | Patent | MARC1 | 50 |
| WO-2014190157-A1 | Table 3. TMPRSS6 single dose screen. | Patent | TMPRSS6 | 47 |
| WO-2021174056-A1 | Table 4. GPR146 Single Dose Screen in Hepa1-6 Cell... | Patent | GPR146 | 44 |
| WO-2016057893-A1 | Table 8. Additional Single Dose Screen in Primary ... | Patent | HAO1 | 44 |
| WO-2015089368-A2 | Table 13. C9 Single dose screen in Primary Mouse H... | Patent | C9 | 44 |
| WO-2020132227-A2 | Table 29. Summary of In Vitro Screening Results fo... | Patent | APP | 43 |
| WO-2016057893-A1 | Table 4a. HAO1 Single Dose Screen in Primary Mouse... | Patent | HAO1 | 42 |
| WO-2022125490-A1 | Table 4. Coagulation Factor X Single Dose Screens ... | Patent | F10 | 41 |
| WO-2021202511-A2 | Table 14. MAPT Single Dose Screens in BE(2)C (huma... | Patent | MAPT | 40 |
| WO-2021222549-A1 | Table 21. In Vitro Single Dose Screen in HepG2 cel... | Patent | CFB | 38 |

Continued on next page

Table S3: Summary of Datapoint Counts per Patent Table (Landscape, 50 chars)

| Patent ID | Table Title | meta_group | Gene | Data (N) |
|---|---|---|---|---|
| WO-2021202511-A2 | Table 15. MAPT Single Dose Screens in NEuro2a (mou... | Patent | MAPT | 38 |
| WO-2021168148-A1 | Table 9. MASP2 Single Dose Screens in Primary Mous... | Patent | MASP2 | 38 |
| WO-2022150260-A1 | Table 4. Single Dose Screen in Hepa1-6 Cells | Patent | C9 | 37 |
| WO-2023056478-A1 | Table 9. Human ANGPTL7 single dose screen in Hepa1... | Patent | ANGPTL7 | 37 |
| WO-2021087036-A1 | Table 31. HTT Single Dose Screens in BE(2)C Cells | Patent | HTT | 36 |
| WO-2021207189-A1 | Table 3: SCN9A in vitro dual luciferase 10nM scree... | Patent | SCN9A | 32 |
| WO-2022178411-A1 | Table 11. In vitro screen of mouse FLCN siRNA in P... | Patent | FLCN | 32 |
| WO-2021087036-A1 | Table 23. HTT Single Dose Screens in PCH Cells | Patent | HTT | 31 |
| WO-2021087036-A1 | Table 22. HTT Single Dose Screens in Hep3B Cells | Patent | HTT | 30 |
| WO-2014190157-A1 | TMPRSS6 single dose screen (10nM) in Hep3B cells w... | Patent | TMPRSS6 | 27 |
| WO-2023003922-A1 | Example 7: In Vitro Single Dose Screening of dsRNA... | Patent | INHBE | 24 |
| WO-2022026531-A1 | Table 4. ATXN2 in vitro screen in Cos-7 (Human Dua... | Patent | ATXN2 | 22 |
| WO-2022178411-A1 | Table 10. In vitro screen of mouse FLCN siRNA in P... | Patent | FLCN | 20 |
| WO-2023003922-A1 | Table 22. Single dose screen for dsRNA agents targ... | Patent | INHBE | 20 |
| WO-2023049871-A2 | Table 10. In vitro Dose Response (\% mRNA Remainin... | Patent | MAPT | 14 |
| WO-2022026531-A1 | Table 7. ATXN2 in vitro screen in Hep3B and BE(2)-... | Patent | ATXN2 | 14 |
| WO-2019089922-A1 | Table 5. C3 single dose screen in Hep3B Cells | Patent | C3 | 12 |
| WO-2021087036-A1 | Table 26. HTT Single Dose Screens in BE(2)C Cells | Patent | HTT | 12 |
| WO-2019089922-A1 | Table 8. C3 single dose screen in Hep3B Cells | Patent | C3 | 12 |
| WO-2021087036-A1 | Table 19. HTT Single Dose Screens in BE(2)C Cells | Patent | HTT | 11 |
| WO-2016057893-A1 | Example 2. In vitro single dose screen in primary ... | Patent | HAO1 | 9 |
| WO-2021022108-A2 | Table 5. CPB2 Single 2 mg/kg Dose Screen in C57BL/... | Patent | CPB2 | 4 |
| WO-2021087036-A1 | Table 13. HTT Single Dose Screens in BE(2)C Cells | Patent | HTT | 4 |
| WO-2021237097-A1 | Table 5. MARC1 in vitro single-dose screen with on... | Patent | MARC1 | 3 |
| WO-2021168148-A1 | Table 10. MASP2 Single Dose Screens in Hep3B Cells | Patent | MASP2 | 2 |

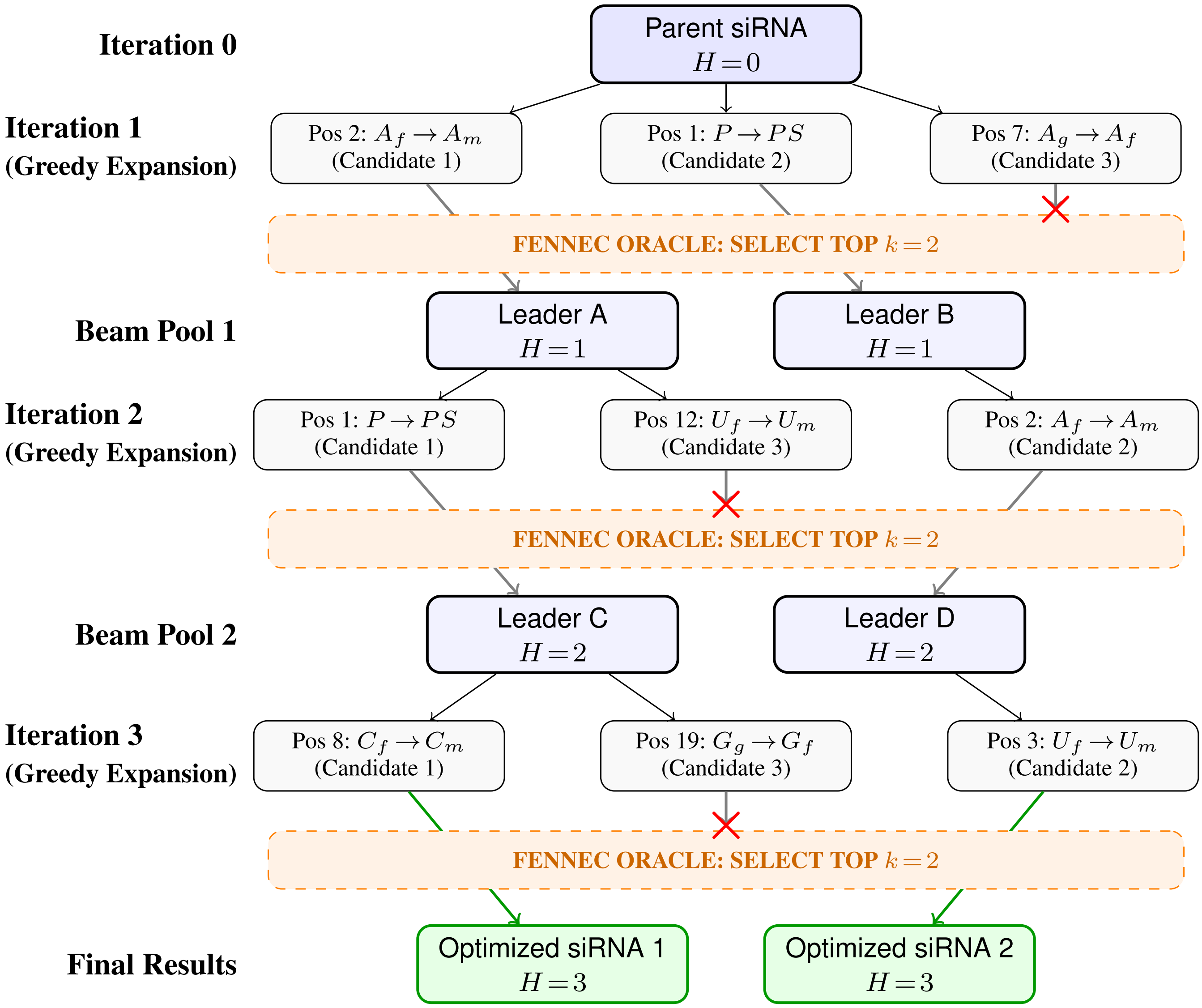


**Figure S9.** Example of the In Silico Sequence Optimization via Greedy Beam Search. K=2 here, while the real design used larger k. Starting from a base parent siRNA ($H=0$), the algorithm iteratively explores single-position chemical edits over three expansion rounds. At each step, 1000s of candidate sequences are evaluated by the FENNEC oracle, which greedily selects the top $k$ performers (blue) to form the next beam pool. Suboptimal variants are discarded (red $\times$). The search concludes with $N$ highly optimized siRNA candidates at a Hamming distance of $H$ (green).

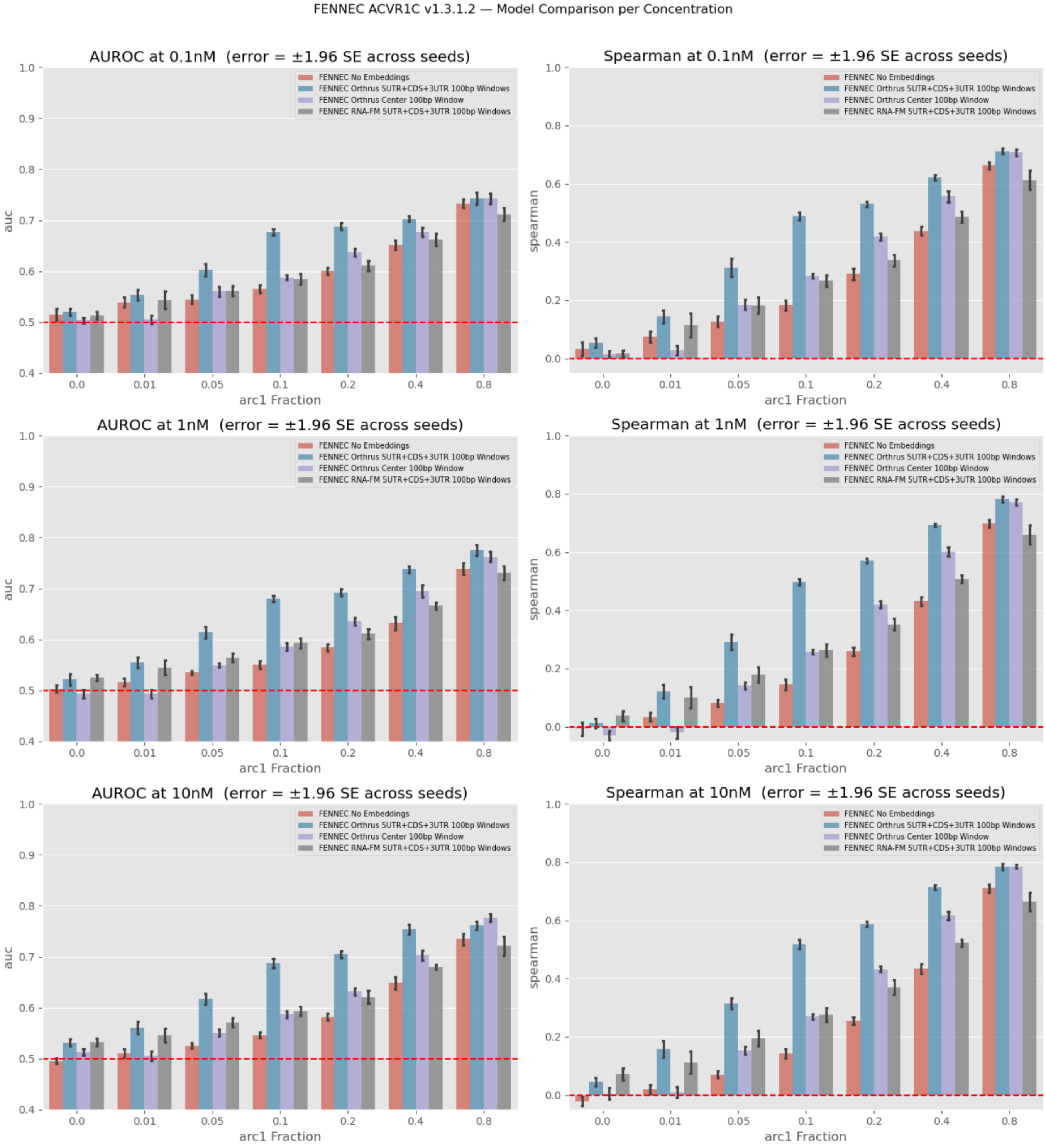


**Figure S10.** FENNEC with Orthrus using different concentrations of data available: 0.1 nM, 1.0 nM, and 10 nM increase in an highly similar manner. Interestingly, the embedding contributions are significantly weaker at 0.1 nM, where the statistical advantage of FM embeddings for ACVR1C is significantly diminished.

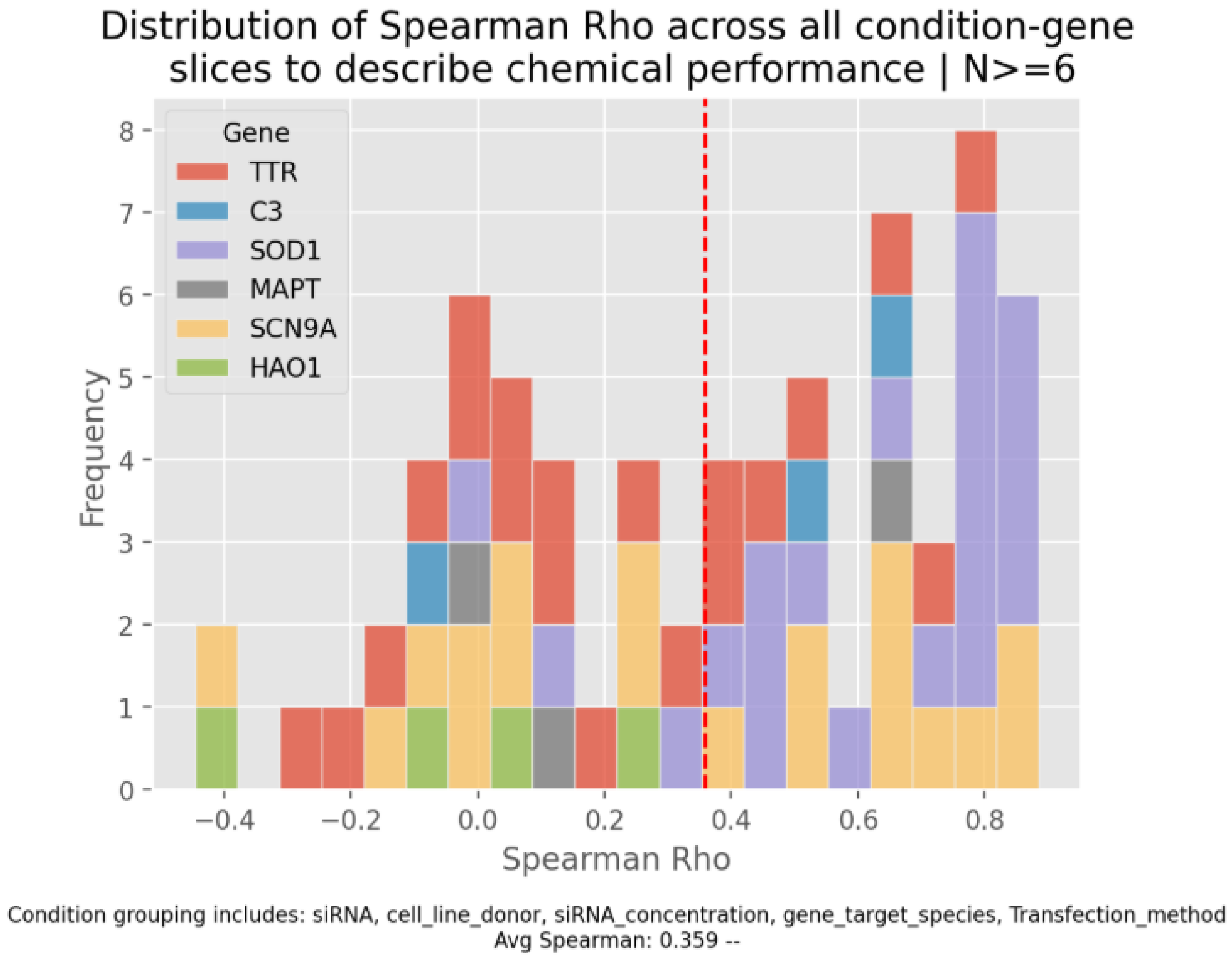


**Figure S11.** CV10 FENNEC 10 model-seed holdout of various grouped conditions to examine the chemical ranking of the model predictions within similar experimental conditions
